# Reconstructing human gene regulatory evolution using deep learning

**DOI:** 10.64898/2026.10.22.760916

**Authors:** Tetsuya Yamada, Xuefei Yuan, Hugo Zeberg, Volker Soltys, Mikel Lana Alberro, Ioannis Sarropoulos, Svante Pääbo, Henrik Kaessmann

## Abstract

Understanding the genetic changes that drove the evolution of human-specific traits is a central goal in biology. Changes in gene expression are thought to be a major source of phenotypic innovations^1,2^, but identifying the underlying changes in cis-regulatory elements (CREs) along the human lineage has been difficult: functional data from great apes are scarce and unobtainable for extinct hominins, and pinpointing causal substitutions requires knowing which sequences determine CRE activity. Here, we used AlphaGenome^3^, a sequence-to-function deep-learning model, to predict the cis-regulatory effects of ∼17 million single-nucleotide substitutions specific to the human lineage and traced their consequences for gene expression evolution across 305 tissues and cell types. The 71,345 predicted cis-regulatory gains lie mostly outside human accelerated regions^4,5^ and arose either through creation of new CREs or, more often, through co-option of CREs ancestrally accessible in other tissues. They are also enriched for distal enhancers and for target gene functions characteristic of the corresponding tissues, suggesting that they contributed to the evolution of tissue-specific biology in humans. Genes linked to more gains were more likely to be upregulated in humans, and in silico reversion to ancestral alleles traced such upregulations to individual substitutions creating transcription factor binding motifs. Comparison with Neandertal and Denisovan genomes revealed that nearly all gains now fixed in humans predate the split from archaic hominins, yet a few modern-human-specific gains contribute substantially to gene upregulation. Our study provides a genome-wide, tissue-resolved map of cis-regulatory changes and their contribution to gene expression evolution along the human lineage.

---

Half a century ago, King and Wilson proposed that the differences between humans and chimpanzees arose mainly from changes in gene regulation rather than in protein sequences^1^. Indeed, most genetic changes (e.g., substitutions) on the human lineage since its split from chimpanzees lie in non-protein-coding regions, where they can alter the activity of cis-regulatory elements (CREs) such as enhancers and promoters. However, it remains largely unknown which of these substitutions changed CRE activity, in which tissues, and at what point in human evolution^6,7^. Functional genomic data from great apes cover only a few tissues and cannot be obtained for extinct hominins. Moreover, even where human-specific changes in CRE activity have been measured, pinpointing the causal substitutions requires knowing which sequence features, such as transcription factor (TF) binding motifs, determine CRE activity.

Sequence-to-function deep-learning models can address both challenges. When trained on sufficiently large and diverse functional genomics data, such as those available for humans, they can learn the sequence features that determine CRE activity and predict non-coding variant effects across tissues and cell types^3,8,9^. Explainable AI methods such as in silico saturation mutagenesis can then pinpoint the nucleotides and motifs underlying each predicted effect^10^. As we and others have shown, because the trans-regulatory environment is largely conserved across mammals, such models can also predict the cis-regulatory potential of orthologous sequences from other species or reconstructed ancestors, allowing cis-regulatory changes between species to be inferred^11,12^. Recent studies have begun to apply this approach to the human lineage^13–15^, but it remains unclear how well the predictions recover measured human-specific cis-regulatory gains and how strongly they are associated with human-specific changes in gene expression.

## Results

### AlphaGenome accurately predicts human-specific cis-regulatory changes

Here, we used AlphaGenome^3^, a state-of-the-art sequence-to-function model, to identify cis-regulatory changes that arose on the human lineage across a broad range of tissues and cell types (Fig. 1a). We focused on 16,958,523 single-nucleotide substitutions for which the human reference genome (GRCh38) carries the derived allele relative to the reconstructed human–chimpanzee ancestral sequence^5^. For each substitution, we predicted chromatin accessibility (DNase I hypersensitivity), a proxy for CRE activity, for the human-derived and ancestral alleles within otherwise identical human sequence, across 305 human tissues and cell types. To make predicted effects comparable across tissues and cell types, we expressed each as a percentile of the predicted effects of common human polymorphisms (minor allele frequency > 0.01) in the same tissue or cell type, which are depleted of variants under strong purifying selection and thus expected to be largely neutral or nearly neutral. We refer to this percentile as the AlphaGenome score: a substitution with a score of 99 has a larger predicted gain in accessibility than 99% of common polymorphisms. This calibration highlights substitutions with unusually large cis-regulatory effects. We refer to predicted human-lineage increases in accessibility as cis-regulatory gains, which can reflect either the emergence of a new CRE or the co-option of an existing CRE into a new tissue context (see below).

**Fig. 1:**
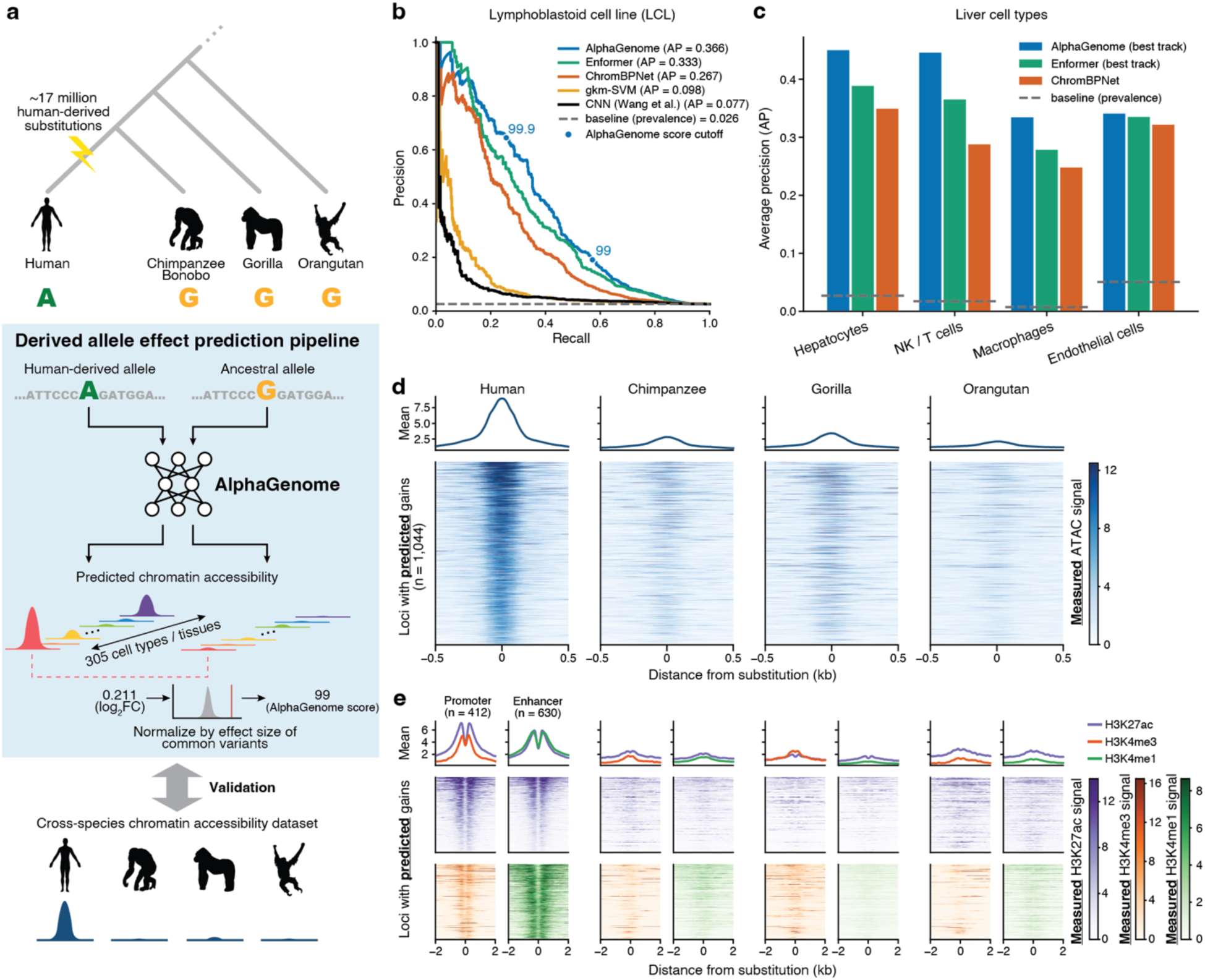
AlphaGenome predicts cis-regulatory elements with human-gained accessibility. **a**, Schematic overview of the workflows for predicting and evaluating cis-regulatory elements (CREs) with human-gained accessibility across cell types and tissues in the human body based on genomes from great apes using AlphaGenome. **b**, Precision–recall curves and average precision (AP) for five sequence models (AlphaGenome, Enformer, ChromBPNet, gkm-SVM, and a classification CNN from Wang et al. 2026^13^) discriminating human-biased differentially accessible ATAC-seq peaks (n = 502) from conserved, non-significant background peaks (n = 18,955) against chimpanzee, gorilla, and orangutan in lymphoblastoid cell lines (LCLs; García-Pérez et al., 2021^16^). Each region is scored by the maximum human-gained-oriented score over substitutions in the same region. Dashed line, prevalence of the positive class against background (0.026). Circles on the AlphaGenome curve mark AlphaGenome score thresholds of 99 and 99.9. **c**, AP for AlphaGenome, Enformer and ChromBPNet in the same task as (**b**) in four liver cell types, for peaks called gained against chimpanzee, bonobo, gorilla and orangutan (hepatocytes, n = 1,038 gained and 37,358 background; NK/T cells, 158 and 8,861; macrophages, 119 and 15,722; endothelial cells, 908 and 17,031). AlphaGenome and Enformer use each cell type’s best-performing tissue or cell type (Supplementary Table 2; Methods). Dashed lines, per-cell-type prevalence. **d**, Mean profile (top) and per-locus heatmap (bottom) of measured chromatin accessibility signal in LCLs from human and each non-human great ape, in ±500 bp windows at the orthologous positions of n = 1,044 predicted accessibility-gained CREs with at least one human-specific substitution with 99.9th percentile AlphaGenome score. Each species’ coverage is scaled by a single multiplicative factor equalizing strong, shared, non-differential peaks to human, so signal heights are comparable across species. Loci ordered by human mean signal. **e**, As in (**d**) (±2.5 kb windows) for histone modifications at the subset of those regions lying in human peaks marking active promoters (H3K4me3+ and H3K27ac+; n = 412) or enhancers (H3K27ac+ and H3K4me1+; n = 630).

We first assessed how accurately AlphaGenome predicts human-specific cis-regulatory changes, using ATAC-seq data from lymphoblastoid cell lines (LCLs) of human, chimpanzee, gorilla, and orangutan^16^, which we reprocessed to identify regions differentially accessible in human relative to all three other species (Methods; Supplementary Table 1). Of 19,457 tested human CREs, 502 had gained accessibility specifically in human. AlphaGenome distinguished these from conserved CREs (average precision 0.366, 14-fold above the 0.026 expected by chance), outperforming all other models tested, including Enformer^17^ and a convolutional neural network classifier used in recent related studies^13,14^, and ChromBPNet^8^ trained directly on the LCL data (Fig. 1b; Supplementary Table 2). AlphaGenome also performed best in predicting human-specific accessibility losses (Extended Data Fig. 1a). In single-nucleus ATAC-seq (snATAC-seq) data of four liver cell types from five great apes^18^, AlphaGenome again ranked first in all four cell types for gains and in three of the four for losses (Fig. 1c; Extended Data Fig. 1b; Supplementary Table 1–2). Most regions with strong predicted changes also showed the corresponding change in measured accessibility, for both gains and losses and across all cell types examined (Extended Data Fig. 1c,d). In LCLs, for example, 82.7% of CREs with a predicted gain (AlphaGenome score ≥ 99.9) were more accessible in human than in each of the other great apes, compared with 31.1% of control CREs with near-zero predicted effects (Fig. 1d). These gains were also accompanied by concordant increases in histone marks of active promoters and enhancers (Fig. 1e). Together, these results show that AlphaGenome accurately predicts human-specific cis-regulatory changes in the tested cell types, supporting its application across the broader range of human tissues and cell types.

### A tissue-resolved map of human-specific cis-regulatory gains

Encouraged by this accuracy, we predicted human-specific cis-regulatory gains genome-wide by combining AlphaGenome scores with human chromatin accessibility data from the ENCODE project^19^ for 305 tissues and cell types. We called a gain when a substitution reached an AlphaGenome score of 99.9 in a given sample and lay within open chromatin in the same sample, a criterion under which precision was 0.65 in the LCL benchmark (25-fold above chance; Fig. 1b). In total, this yielded 128,646 regions with predicted gains (Supplementary Table 3). Because cell lines and cultured non-immune primary cells clustered by culture state rather than by tissue of origin (Extended Data Fig. 2a), we focused subsequent analyses on 144 primary tissues, in vitro-differentiated cell types and immune cells, in which 71,345 regions had predicted gains (Supplementary Table 4; Methods). Clustering these tissues and cell types by their predicted gains, considering each region only where it is accessible, yielded 25 tissue programs spanning diverse organ systems. Each region was assigned to the program in which its predicted gain was strongest, except for regions with gains in at least 10% of tissues and cell types, which formed a broad class (Fig. 2a; Extended Data Fig. 2b,c). In each of the 25 programs, gains were significantly associated with the creation of binding motif instances for TFs characteristic of the constituent tissues, identified from deep-learning-based motif annotations (ENCODE-GRAMMAR^20^; Methods). Examples include ETS-domain TFs in immune cells, HNF TFs in liver and intestine, and SOX family TFs in the central nervous system. In contrast, gains in the broad class were associated with the creation of CTCF motif instances (Fig. 2a; Supplementary Table 5), which may give rise to human-specific chromatin domain boundaries acting across cell types^21^. Similarly, genes near the gained regions were enriched for functions characteristic of the corresponding tissues in 22 of 25 programs (Fig. 2a; Supplementary Table 6), suggesting that the predicted regulatory gains contributed to the evolution of tissue-specific biology in humans.

**Fig. 2:**
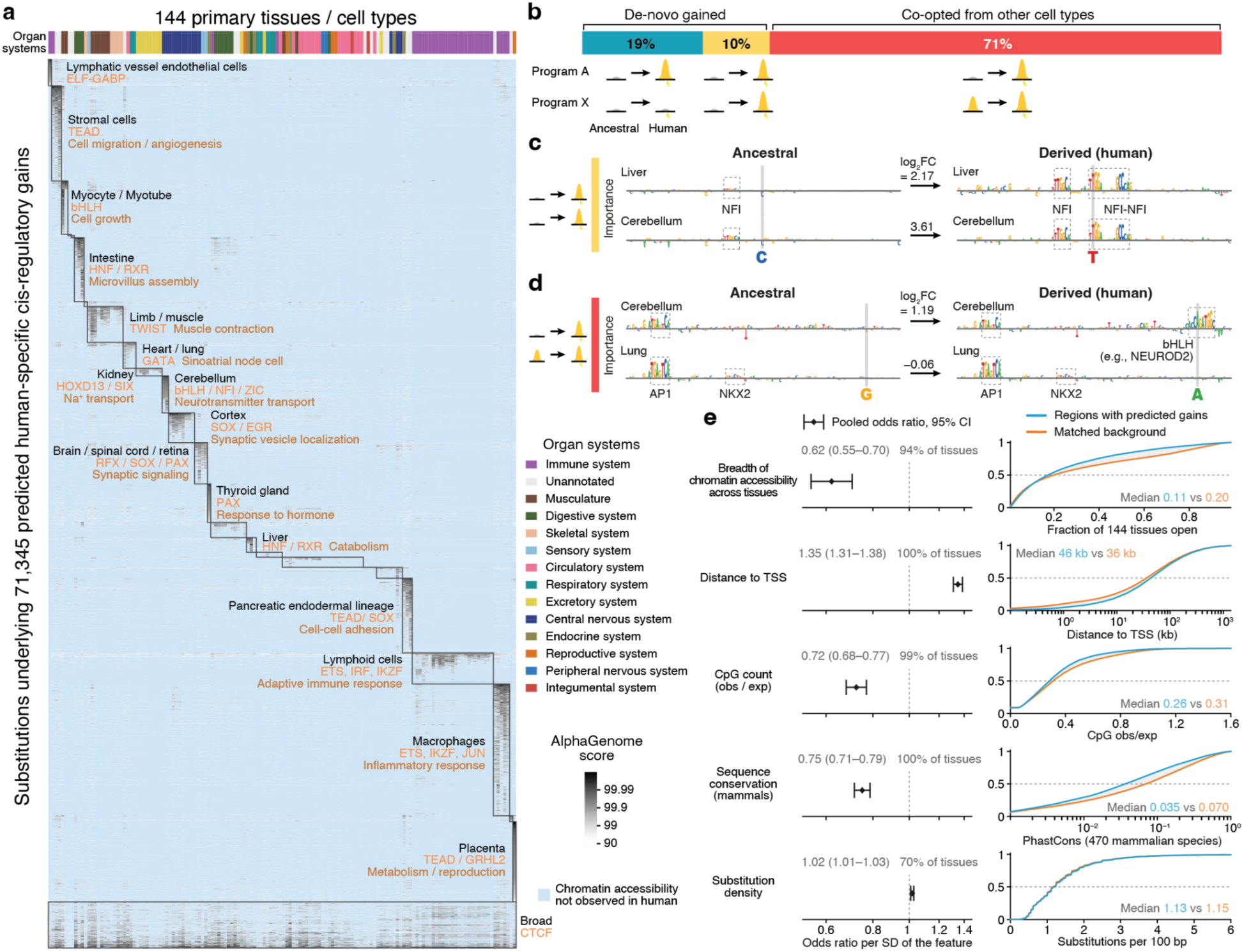
Global patterns of predicted CREs with human-gained accessibility. **a**, Clustered heatmap of AlphaGenome scores for predicted accessibility-gained CREs across 144 tissues and cell types. Rows are a random sample of 10,000 of the 71,345 elements with an AlphaGenome score ≥ 99.9 in at least one tissue or cell type, ordered by their assigned program. Columns are tissues and cell types, clustered hierarchically into 25 programs. Color indicates the AlphaGenome score of the strongest substitution within each element, on a log scale. Scores are shown only where the element overlaps a DNase-seq peak in the matched ENCODE biosample. Major clusters are annotated by their constituent tissues, by TF motifs enriched among the human-lineage substitutions, and by enriched biological functions of nearby genes. **b**, Mode of accessibility gain for the 68,031 predicted accessibility-gained CREs assigned to one of the 25 tissue programs. Accessibility is an ENCODE DNase-seq peak in human at the substitution, and gains are called at the 99.9th percentile of the AlphaGenome score. **c,d**, Examples of a de novo gain shared across programs (**c**) and of co-option (**d**), at chr17:72,589,117–72,589,218 and chr9:27,655,049–27,655,150, respectively. Contribution of each base to the AlphaGenome accessibility prediction, from in silico saturation mutagenesis over ±50 bp around the human-specific substitution, is shown for the reconstructed ancestral sequence (left) and the human sequence (right), on cerebellar cortex (top) and left lobe of liver (bottom) in (**c**), and on cerebellum (top) and upper lobe of left lung (bottom) in (**d**). High-importance matches to known TF binding motifs and the substitution are annotated. **e**, (Left) Random-effects meta-analysis of per-tissue multivariable logistic regressions (142 tissues with at least 50 predicted human-gained calls) of whether a human-lineage substitution is called gained. Points are the average effect across tissues, with 95% confidence intervals from a random-effects model that allows the true effect to differ between tissues. Beside each row is the fraction of tissues whose own estimate points in the same direction as the average. (Right) Empirical cumulative distribution of each covariate for accessibility-gained CREs and for background CREs reweighted by inverse probability of being gained.

As introduced above, a cis-regulatory gain can arise either de novo or through co-option of a CRE that was already accessible in other tissues^22,23^. The relative contribution of these two modes has been difficult to assess. By combining AlphaGenome scores with measured chromatin accessibility in human, we classified each predicted gain by whether its CRE is also accessible in other tissue programs and, if so, whether the substitution is predicted to account for that accessibility as well (Methods). Of the 68,031 regions with gains assigned to one of the 25 tissue programs (that is, excluding the broad class), 29% were consistent with de novo emergence: the CRE was accessible either only in the program in which the gain was called (19%) or also in other programs in which the substitution was predicted to increase accessibility (10%) (Fig. 2b). In the latter case, the substitution tended to create a motif instance for TFs active across tissue programs, such as AP-1 (FOS–JUN) and NFI^24^ (Fig. 2c; Extended Data Fig. 3a). The remaining 71% were accessible in at least one other program without a comparable predicted gain, suggesting that the CRE was ancestrally accessible and that the substitution drove its co-option into a new program (Fig. 2b), in line with the predominance of co-option reported for cell types of the developing brain^25^. These co-opted CREs tended to retain existing motif instances for broadly active TFs, such as NRF and AP-1, while the substitution added a motif instance for a tissue-specific TF, such as PAX, HNF or NEUROD^11,24^ (Fig. 2d; Extended Data Fig. 3b,c). AP-1 motif instances thus contributed both to de novo gains shared across tissue programs and to the permissive background of co-opted CREs. Overall, human-specific cis-regulatory gains mostly arose at elements already used as CREs in other tissues, where existing motif instances for broadly active TFs may provide a permissive starting point for tissue-specific gains^22,23^.

We next characterized the regions with predicted cis-regulatory gains, comparing them with other accessible regions in the same tissues in a joint model, so that each feature was assessed independently of the others and of potential confounders, such as GC content and accessibility signal strength (Methods). These regions were accessible in fewer tissues, more often distal to transcription start sites (TSSs), CpG-poor, and less constrained across mammals (Fig. 2e), consistent with an enrichment for putative distal enhancers: 71.0% overlapped ENCODE distal enhancer-like elements, compared with 57.2% of accessibility-matched control regions (odds ratio [OR] = 1.83; *P* < 10^−300^, Fisher’s exact test; Extended Data Fig. 4a,b). By contrast, their density of human-lineage substitutions was nearly identical to that of controls (Fig. 2e).

These properties set them apart from “human accelerated regions” (HARs)^4^ and “human ancestor quickly evolved regions” (HAQERs)^5^, the non-coding sequences that have attracted much attention as candidate contributors to human evolution. Both are defined by an elevated substitution rate on the human lineage, HARs additionally by prior conservation. Neither criterion applies to the regions identified here, and the overlap was small: only 157 of the 71,345 regions overlapped a HAR (5.8% of the 2,698 HARs), and 76 overlapped a HAQER (4.8% of the 1,581 HAQERs). Thus, identifying regions by the predicted effects of individual substitutions recovers a largely different set of non-coding regions from approaches based on substitution rates. Such rates can be elevated by processes unrelated to regulatory activity, such as GC-biased gene conversion, which affects HARs and HAQERs more strongly than the regions identified here (Extended Data Fig. 5a,b), as well as relaxed constraint or high local mutation rates^5,26^.

Human-specific enhancers have been reported to cluster in the genome rather than being randomly distributed^27^. Our data confirmed this across LCLs and the four liver cell types. Within cell types, pairs of human-specific gains, and likewise pairs of losses, occurred within 3–10 kb of each other at about twice the rate expected by chance. Predicted gains also clustered within tissue programs, but more weakly than measured gains (Extended Data Fig. 6a). Prediction error alone did not account for this: errors distributed at random relative to the measured gains yielded more close pairs than observed (empirical one-sided *P* = 0.038; Extended Data Fig. 6b), suggesting that clustered gains are missed more often than isolated ones. Consistent with this, measured gains with nearby gains were less often recovered by AlphaGenome, from 45.0% for isolated gains to 28.5% for those with two or more neighbors within 30 kb (Extended Data Fig. 6c). Those missed within clusters showed smaller accessibility changes than their recovered neighbors (Extended Data Fig. 6d). Such gains may thus reflect accessibility changes not driven by substitutions within the affected element itself, for instance through coordinated regulation with nearby CREs^28^. Together, these results show that human-specific cis-regulatory gains tend to occur near each other within a given tissue, and that AlphaGenome predictions mainly capture the subset driven by the element’s own substitutions (Extended Data Fig. 6e).

### Linking cis-regulatory gains to gene expression evolution

We next asked whether these cis-regulatory gains contribute to human-specific changes in gene expression. Because gains tend to occur near each other, a single gene may be linked to several of them, which could jointly upregulate its expression in humans. To test this, we linked CREs to their target genes using ENCODE-rE2G^29^, a high-accuracy enhancer–gene map, in the tissue or cell type matching each expression dataset (Supplementary Table 7). For each gene, we summed the link scores of all its CREs with predicted gains, a quantity we call the gained rE2G burden (Fig. 3a; Methods). Because this burden aggregates over many regions and is therefore less sensitive to individual false positives, we favored sensitivity over precision and called gains at an AlphaGenome score of 99 instead of 99.9 (precision 0.19 in the LCL benchmark, seven-fold above chance; Fig. 1b). We then compared the gained rE2G burden with expression differences between human and other great apes in ten published transcriptome datasets^11,18,30^, covering four liver cell types, cerebellar granule cells, and five organs.

**Fig. 3:**
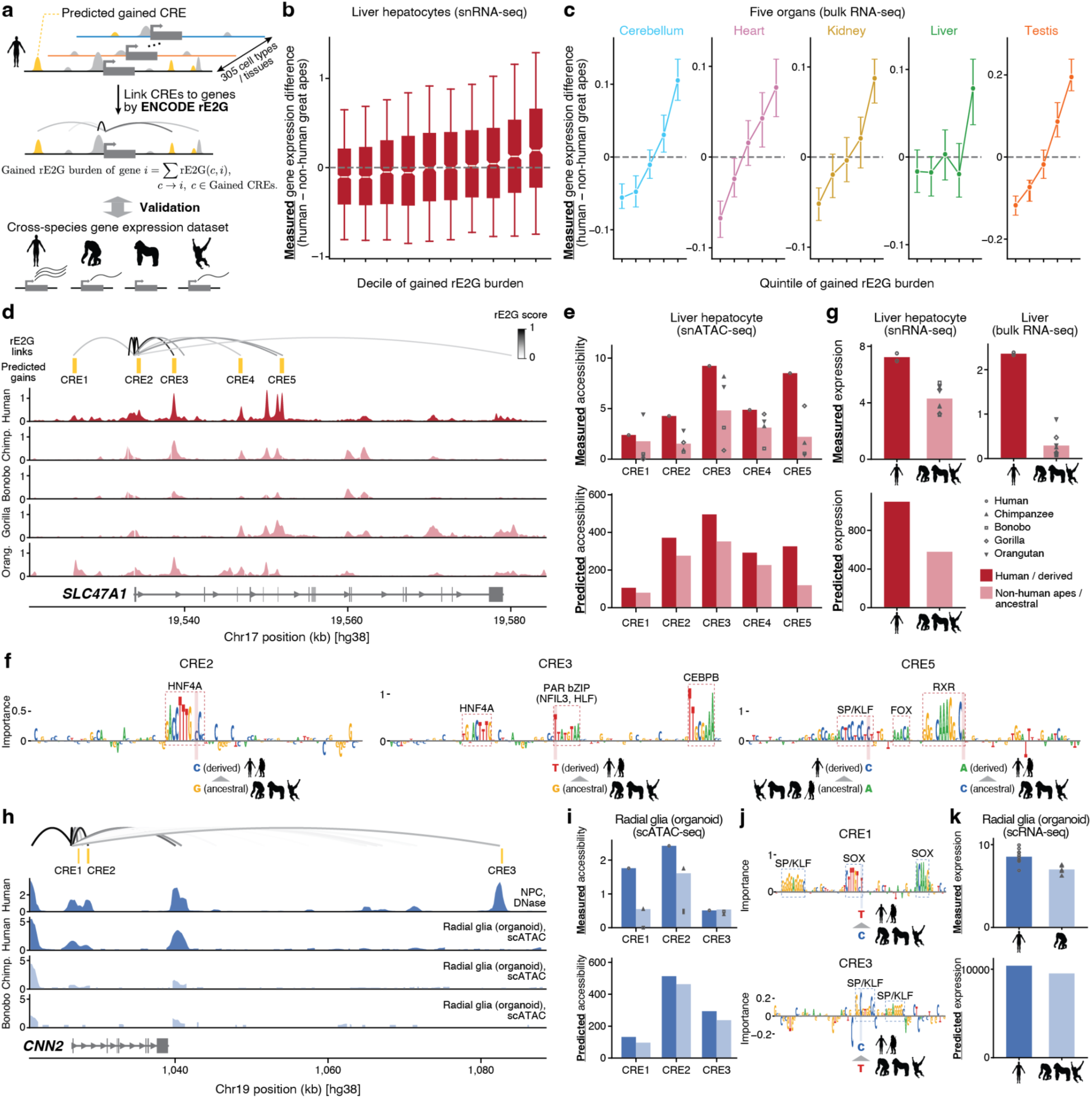
Predicted accessibility-gained CREs link human-specific substitutions to gene expression evolution across tissues. **a**, Schematic overview of the workflow for aggregating predicted accessibility-gained CREs into a per-gene burden using ENCODE-rE2G regulatory element–gene links from matching tissues. **b**, Difference in gene expression levels between human and four non-human great apes (chimpanzee, bonobo, gorilla, orangutan) in hepatocytes, from the Yuan et al., 2025^18^ single-nucleus RNA-seq (snRNA-seq) pseudobulk, against deciles of rE2G burden of predicted gained CREs. Gained CREs are hepatocyte snATAC peaks containing at least one substitution called human-gained by AlphaGenome scored on those peaks for the right lobe of liver. rE2G burden is the sum of ENCODE-rE2G link scores from the left lobe of liver over a gene’s gained CREs at 99th percentile AlphaGenome score. Genes were retained above a symmetric expression floor on the average of the human and mean-ape values, leaving 13,322 genes, 1,332–1,333 per decile. Expression differences are median-centered within the dataset. Boxes show the interquartile range, whiskers the 10th–90th percentiles, and notches the 95% confidence interval of the median. **c**, Median difference in expression between human and four non-human great apes (chimpanzee, bonobo, gorilla, and orangutan) with percentile bootstrap 95% confidence intervals, against quintiles of rE2G burden of predicted gained CREs, in five organs from the Brawand et al., 2011^30^ bulk RNA-seq. Gained CREs and rE2G links come from the DNase biosample matched to each tissue. Expression differences are median-centered within each tissue. **d**, The *SLC47A1* locus. From top: ENCODE-rE2G links to the *SLC47A1* TSS colored by link score; the five predicted accessibility-gained CREs; measured accessibility in human; measured accessibility in non-human great apes, at the orthologous positions and scaled to human by anchor peaks; and the gene model. **e**, For each of the five CREs in (**d**): measured ATAC coverage in human against the great-ape mean (top), and AlphaGenome-predicted accessibility (right lobe of liver) for the human sequence against the same sequence carrying the reconstructed ancestral states (bottom). **f**, Contribution of each base of the human sequence to the AlphaGenome accessibility prediction, from in silico saturation mutagenesis over ±50 bp. Shown for three of the five CREs in (**d**). High-importance matches to known TF binding motifs and the human-specific substitutions are annotated. **g**, (Top) Measured expression of *SLC47A1* in human and four great apes, in hepatocytes by snRNA-seq (left; points are donor pseudobulks) and in whole liver by bulk RNA-seq (right). (Bottom) AlphaGenome-predicted *SLC47A1* expression (hepatocyte, total RNA-seq) for the human sequence and for the same sequence with all five gained CREs reverted to their ancestral states simultaneously. **h**, The *CNN2* locus. From top: ENCODE-rE2G links to the *CNN2* TSS colored by link score; the three predicted accessibility-gained CREs; measured accessibility from DNase-seq in human neural progenitor cells (NPCs); measured accessibility from single-cell ATAC-seq of radial glial cell pseudobulk in cerebral organoids of human, chimpanzee, and bonobo from Kanton et al., 2019^32^; and the gene model. **i**, For each of the three CREs in (**h**), measured ATAC coverage in human against the great-ape mean (top) and AlphaGenome-predicted accessibility (NPCs) for the human sequence against the same sequence carrying the reconstructed ancestral states (bottom). **j**, As in (**f**), for two of the three CREs in (**h**) over ±40 bp. **k**, (Top) Measured expression of *CNN2* in human and chimpanzee in radial glia pseudobulk in cerebral organoids by scRNA-seq from Kanton et al., 2019^32^. (Bottom) AlphaGenome-predicted *CNN2* expression (NPCs) for the human sequence and for the same sequence with all three gained CREs reverted to their ancestral states simultaneously.

Across all ten datasets, genes with a higher gained rE2G burden, that is, with more gains strongly linked to them, were more likely to be upregulated in humans relative to other great apes. In hepatocytes^18^, for example, the human–ape expression difference increased steadily with gained rE2G burden (Fig. 3b), and the fraction of human-upregulated genes rose from 41% to 62% (OR = 2.4; *P* = 3 × 10^−29^, Fisher’s exact test). This association remained significant after controlling for the burden of the gene’s other CREs, its dosage sensitivity, and its ancestral expression level (*P* = 4 × 10^−15^, Wald test), indicating that it is specific to the gained CREs. The same gradient appeared in the other three liver cell types^18^ (*P* = 6 × 10^−5^ to 2 × 10^−24^; Extended Data Fig. 7a), in cerebellar granule cells compared between human and bonobo^11^ (*P* = 5 × 10^−47^; Extended Data Fig. 7b), and in five organs profiled by bulk RNA-seq^30^ which we requantified on a consensus gene annotation across species (*P* = 1 × 10^−4^ to 2 × 10^−45^; Fig. 3c; Methods). By contrast, burdens computed from HARs, HAQERs, or human-lineage-gained CREs from two recent related studies^13,14^ were significantly associated with human upregulation in only 0, 1, 3, and 3 of the 10 datasets, respectively (Extended Data Fig. 7c), although individual accelerated regions were shown to contribute to human-specific gene upregulations^31^. Therefore, predicted cis-regulatory gains identify not only human-specific changes in CRE activity but also genes likely to have changed expression in humans, without requiring functional data from other great apes.

Genes with a high gained rE2G burden are promising candidates for adaptive changes in expression. Because each is linked to several independently predicted regulatory gains, expression differences at these genes are more likely to reflect substitutions acting in cis than trans-regulatory differences, variation among sampled individuals or technical artifacts. Moreover, multiple gains acting in the same direction on a single gene are consistent with directional selection. In agreement with this notion, genes in the top decile of gained rE2G burden shared properties reported for genes with signatures of positive selection on their protein-coding sequences^33,34^: compared with other genes, they were more tissue-specific, more often encoded secreted, extracellular-matrix, or membrane proteins rather than nuclear ones, and were less often among the most dosage-sensitive genes (Extended Data Fig. 8; Methods). This suggests that adaptive changes in gene regulation and in protein sequences target similar classes of genes.

To identify the strongest candidates for adaptive expression changes, we focused on genes with both a high gained rE2G burden and measured human-specific upregulation (Supplementary Table 8). We then traced their upregulation to individual regions with predicted cis-regulatory gains and to the underlying substitutions. We further used AlphaGenome to predict the expression consequences of these substitutions directly, by reverting them to their ancestral alleles in silico (Methods). One candidate is *SLC47A1*, which encodes MATE1, a key transporter in the excretion of drugs, toxins and waste products via the liver and kidney^35,36^. *SLC47A1* is in the top 0.5% of genes by gained rE2G burden in hepatocytes and is linked to five regions with predicted gains within 26 kb of its promoter (Fig. 3d). All five regions were more accessible in human hepatocytes than in any of the four other great apes, and AlphaGenome recapitulated this from sequence alone: reverting the human-lineage substitutions to their ancestral states reduced predicted accessibility at each region (Fig. 3e), with effect sizes tracking the measured differences. In three of the five regions, explainable AI analysis based on in silico saturation mutagenesis showed that the substitution created a motif instance for a hepatocyte-specific TF (Fig. 3f). Consistent with these liver-specific regulatory gains, *SLC47A1* was upregulated in humans relative to other great apes in hepatocytes and bulk liver (Fig. 3g), but not in kidney, where it is expressed in all great apes and no gains were predicted (Extended Data Fig. 9a,b). Simultaneously reverting the substitutions in all five regions to their ancestral alleles reduced its predicted expression by 60% (Fig. 3g), suggesting that these gains jointly account for much of its upregulation in humans.

Neural progenitor cells (NPCs) are of particular interest, because their increased proliferative capacity is thought to underlie the expansion of the human neocortex^37^. Among the candidates in this cell type is *CNN2*, which encodes calponin-2, an actin-binding protein that regulates cell adhesion, migration, and proliferation^38^. *CNN2* is in the top 2% of genes by gained rE2G burden in NPCs and is linked to three regions with predicted gains within 60 kb of its promoter (Fig. 3h). Two of these regions were also more accessible in human than in chimpanzee and bonobo radial glia from cerebral organoids^32^ (Fig. 3h,i), and the human-lineage substitutions created motif instances for SOX and SP/KLF (Fig. 3j). *CNN2* is upregulated in human relative to chimpanzee radial glia in cerebral organoids^32,39^, and reverting the substitutions in all three regions to their ancestral states reduced its predicted expression by 7% (Fig. 3k), suggesting that these cis-regulatory gains account for part of this upregulation.

Further candidates include *ZP2* in cerebellar granule cells, previously shown to have gained expression in adult human granule cells and to be involved in synapse development and neuronal activity^11,40,41^, *ADAMTS17* in NK/T cells, *CD226* in macrophages, and *JCAD* in endothelial cells of the liver (Extended Data Fig. 10a–d), as well as *CSDC2* in heart, *ALS2CL* in kidney, and *MRC2* in testis (Extended Data Fig. 11a–c). Together, these examples show how human-specific upregulation of genes can be traced to predicted cis-regulatory gains and to the underlying substitutions, highlighting candidate targets of adaptive changes in gene expression during human evolution.

### Identifying modern-human-specific regulatory and expression changes

Our approach, which only requires DNA sequence as input, provides a unique opportunity to determine when cis-regulatory gains and associated expression changes arose in humans relative to the divergence from extinct hominins, for which functional genomic data cannot be obtained. Using seven high-coverage archaic genomes, from five Neandertals^42–45^ and two Denisovans^46,47^, we dated human-lineage substitutions at sites where these genomes could be genotyped. A substitution was classified as modern-human-specific when all genotyped archaic genomes carried the ancestral allele and as shared with archaic hominins when all genotyped archaic genomes carried the derived allele (Fig. 4a; Methods). Each region took the age of its youngest substitution with a gain. Of the 69,803 autosomal regions with predicted gains, 4,902 (7.0%) were modern-human-specific, 50,975 (73.0%) were shared with archaic hominins, and the remainder could not be resolved (Table 1; Supplementary Table 4). Among regions whose age could be inferred, 8.8% were modern-human-specific, significantly below the 10.0% observed across all human-lineage substitutions (*P* = 6 × 10^−55^, Fisher’s exact test) and in line with the ∼9.2% expected from population divergence times alone (∼0.6 versus ∼6.5 million years since the splits from archaic hominins and from chimpanzees, respectively)^48,49^. The substitutions underlying predicted gains are thus enriched among those that arose before the split from archaic hominins.

**Fig. 4:**
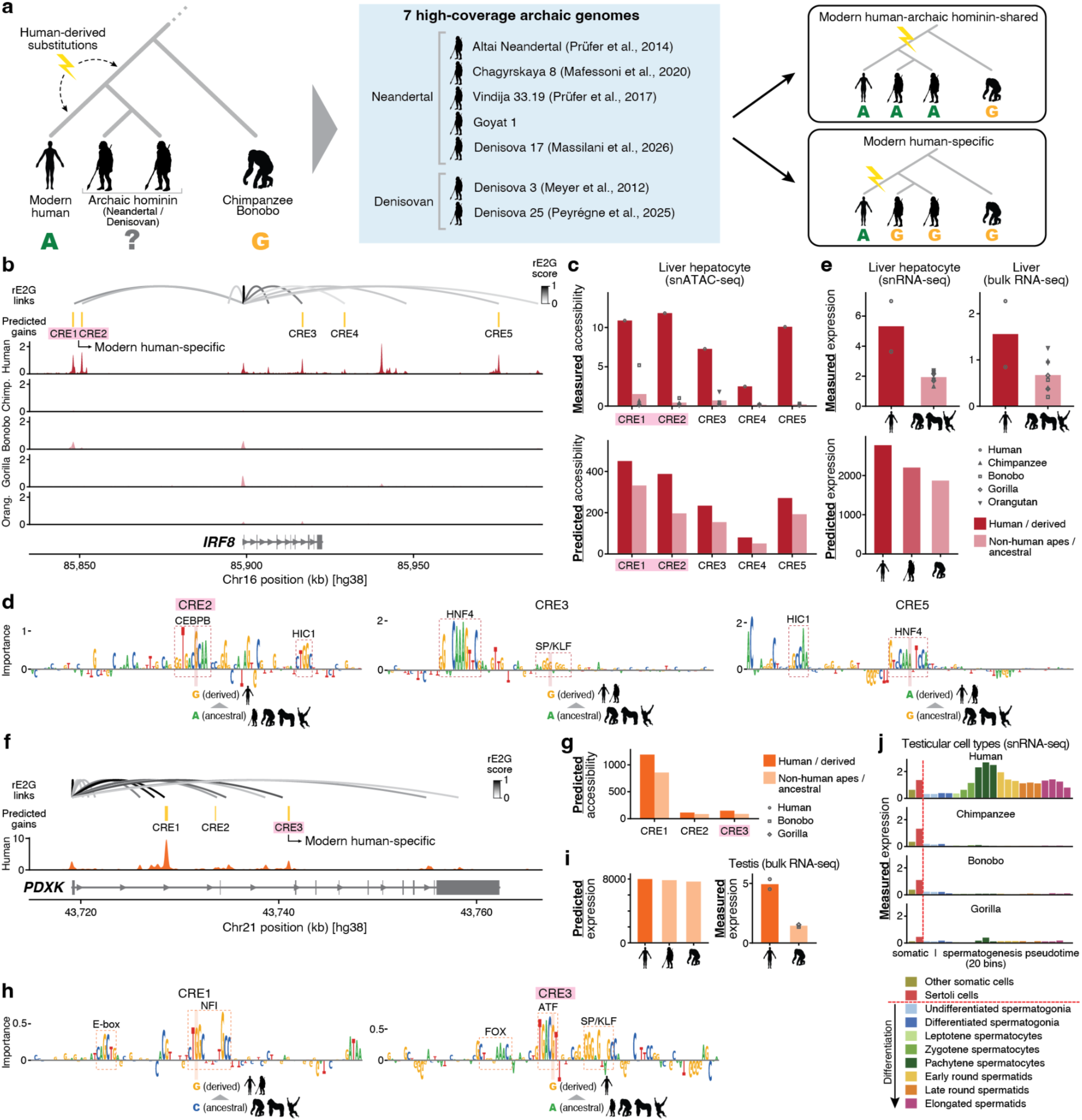
CREs with modern-human-gained accessibility predicted to contribute to gene expression evolution. **a**, Schematic overview of the workflows for identifying modern-human-specific cis-regulatory gains. **b**, The *IRF8* locus. From top: ENCODE-rE2G links to the *IRF8* TSS colored by link score; the five predicted accessibility-gained CREs, with those specific to modern humans highlighted; measured accessibility in human; measured accessibility in non-human great apes, at the orthologous positions and scaled to human by anchor peaks; and the gene model. **c**, For each of the five CREs in (**b**): measured ATAC coverage in human against the great-ape mean (top), and AlphaGenome-predicted accessibility (right lobe of liver) for the human sequence against the same sequence carrying the reconstructed ancestral states (bottom). **d**, Contribution of each base of the human sequence to the AlphaGenome accessibility prediction, from in silico saturation mutagenesis over ±50 bp. Shown for three of the five CREs in (**b**). High-importance matches to known TF binding motifs and the human-specific substitutions are annotated. **e**, (Top) Measured expression of *IRF8* in human and four great apes, in hepatocytes by snRNA-seq (left; points are donor pseudobulks) and in whole liver by bulk RNA-seq (right). (Bottom) AlphaGenome-predicted *IRF8* expression (hepatocyte, total RNA-seq) for the human sequence and for the same sequence with all five gained CREs reverted to their ancestral states simultaneously. **f**, The PDXK locus. From top: ENCODE-rE2G links to the *PDXK* TSS colored by link score; the three predicted accessibility-gained CREs, with those specific to modern humans highlighted; measured accessibility in human; and the gene model. **g**, For each of the three CREs in (**f**), AlphaGenome-predicted accessibility (testis) for the human sequence against the same sequence carrying the reconstructed ancestral states. **h**, As in (**d**), for two of the three CREs in (**f**). **i**, (Left) AlphaGenome-predicted *PDXK* expression (testis, total RNA-seq) for the human sequence and for the same sequence with all three gained CREs reverted to their ancestral states simultaneously. (Right) Measured expression of *PDXK* in human and two great apes (bonobo and gorilla) in whole testis by bulk RNA-seq. **j**, Measured expression of *PDXK* in testicular cell types in human and three great apes, based on snRNA-seq data from Murat et al., 2023^54^.

**Table 1:** Regions with predicted human-specific cis-regulatory gains on autosomes partitioned by allelic state in seven archaic hominins and by derived allele frequency (DAF) in African populations from the 1000 Genomes Project.

|  | Fixed | Derived Allele Frequency (DAF) bins |  |  |  | Total |
| --- | --- | --- | --- | --- | --- | --- |
|  |  | 0.99–1 | 0.90–0.99 | 0.50–0.90 | <0.50 |  |
| Modern-human-specific | 172 | 102 | 297 | 1,335 | 2,996 | 4,902 |
| Shared with archaic hominins | 47,550 | 894 | 967 | 1,198 | 366 | 50,975 |
| Unresolved | 10,081 | 491 | 781 | 1,518 | 1,055 | 13,926 |
| All gained CREs | 57,803 | 1,487 | 2,045 | 4,051 | 4,417 | 69,803 |

Because substitutions were defined from the reference genome, they need not be fixed in present-day humans. This applies especially to modern-human-specific substitutions, which have had far less time to spread through the population than those shared with archaic hominins and are therefore expected to be fixed less often and to occur more often at lower derived allele frequencies (DAFs). Yet fixed or nearly fixed changes are of particular interest, as they are shared by virtually all present-day humans and could thus contribute to traits that distinguish modern humans from archaic hominins^6^. We therefore examined DAFs in the five African populations of the 1000 Genomes Project^50^, which harbor most present-day human genetic variation and carry little archaic ancestry. Only 172 of the 4,902 modern-human-specific gains (3.5%) were fixed in these populations, compared with 47,550 of the 50,975 gains shared with archaic hominins (93.3%; Table 1; Supplementary Table 4). Consequently, almost all cis-regulatory gains now fixed in present-day humans (99.6%) arose before the split between modern humans and archaic hominins. Moreover, modern-human-specific substitutions predicted to cause gains more often reached a DAF of at least 0.99 than modern-human-specific substitutions genome-wide (5.8% versus 4.2%; *P* = 2 × 10^−7^, Fisher’s exact test). Neither this excess nor the excess of gains shared with archaic hominins was explained by GC-biased gene conversion, which affects HARs and HAQERs more strongly (Extended Data Fig. 12a,b; Methods). These results are therefore consistent with gain-causing substitutions having reached high frequency or fixation more rapidly than substitutions genome-wide, as expected under positive selection acting on a subset of them. In line with this, fixed modern-human-specific substitutions predicted to cause regulatory gains fell within candidate regions of ancient selective sweeps on the modern human lineage^51^ more often than other fixed modern-human-specific substitutions in open chromatin (14.4% versus 8.2%; *P* = 0.006, Fisher’s exact test).

Although rare, modern-human-specific cis-regulatory gains can contribute substantially to human-specific gene expression increase. One example is *IRF8* in hepatocytes. *IRF8* encodes a TF best known for its role in immune cell differentiation and function, but genetic variation at the *IRF8* locus has also been associated with circulating cholesterol levels^52^, suggesting a possible role in hepatic lipid metabolism. *IRF8* is in the top 1% of genes by gained rE2G burden in hepatocytes and is strongly linked to five regions with predicted gains (Fig. 4b,c), two of which (CRE1 and CRE2) are driven by modern-human-specific substitutions. The substitution in CRE2 is nearly fixed (DAF 0.98 in African populations of the 1000 Genomes Project^50^) and creates a motif instance for the hepatocyte TF CEBPB (Fig. 4d). *IRF8* is also in the top 0.5% of genes with human-specific expression increases in hepatocytes and bulk liver. Reverting the substitutions in all five regions to their ancestral states in silico recapitulated this expression increase (Fig. 4e). Our analyses further indicate that *IRF8* expression increased in two steps along the human lineage, first through substitutions that arose before the divergence from archaic hominins and then through modern-human-specific ones, which account for ∼47% of the predicted change (Fig. 4e).

Another example is *PDXK* in testis, which encodes pyridoxal kinase, the enzyme that activates vitamin B6^53^. *PDXK* is in the top 0.5% of genes by gained rE2G burden in testis and is linked to three regions with predicted gains within 22 kb of its promoter (Fig. 4f,g). One of these (CRE3) is driven by a modern-human-specific substitution that is fixed in all five African populations and creates a motif instance for bZIP TFs of the ATF/CREB family (Fig. 4h). *PDXK* is in the top 0.2% of genes by human-specific expression increase in testis (Fig. 4i). Reverting the substitutions in the three regions in silico recapitulated this increase only weakly, with the modern-human-specific substitution accounting for ∼42% of the predicted change (Fig. 4i). A cross-primate single-nucleus RNA-seq atlas of testis^54^ suggests a potential reason: *PDXK* is expressed mainly in Sertoli cells in non-human great apes, whereas in humans it is additionally, and much more highly, expressed in pachytene spermatocytes (Fig. 4j), a cell type whose regulatory landscape bulk-testis accessibility data and the corresponding predictions may only partially capture. Together, these results indicate that, although most cis-regulatory gains along the human lineage arose before the split from archaic hominins, a small number of more recent, modern-human-specific gains can have substantial effects on gene expression and may have contributed to traits that distinguish modern humans from archaic hominins.

## Discussion

In summary, we predicted the cis-regulatory effects of substitutions that arose on the human lineage, genome-wide and across tissues, together with their contributions to gene expression evolution. AlphaGenome accurately predicted measured human-specific changes in accessibility, outperforming models used in recent related studies^13,14^, and allowed us to pinpoint the substitutions and motifs underlying individual gains. The predicted gains are enriched for putative distal enhancers and mostly arose through co-option of CREs already accessible in other tissues, rather than through the emergence of new CREs. They are almost entirely distinct from HARs^4^ and HAQERs^5^, and were far more consistently associated with human-specific upregulation of linked genes than these and other recently proposed candidate regions^13,14^, allowing us to trace such upregulation to individual regulatory gains and the underlying substitutions. Combining the predictions with archaic hominin genomes further showed that nearly all gains now fixed in humans predate the split from Neandertals and Denisovans, and that gain-causing substitutions show patterns consistent with positive selection on a subset of them. Although far fewer, modern-human-specific regulatory gains also contribute substantially to gene upregulation. To facilitate follow-up studies, we provide all predictions and analyses through an interactive web application (https://apps.kaessmannlab.org/human_specific_cre), where users can explore predicted human-specific cis-regulatory changes and their aggregated effects on gene expression.

A limitation of our study is the sensitivity of the predictions. Although AlphaGenome is the state-of-the-art model and outperformed all others tested, it recovers roughly a quarter to half of measured human-specific accessibility changes in our LCL benchmark, depending on the threshold. It misses clustered gains more often than isolated ones, suggesting that some human-specific changes are driven by sequence outside the affected element, for instance through coordinated regulation of nearby CREs^28^. Functional genomic data from other species therefore remain necessary for a complete view of cis-regulatory change, although the sensitivity of sequence-based predictions is bound to increase as models improve. The sequence-based approach nonetheless offers what measured data cannot: it isolates cis-regulatory effects from trans-regulatory differences and variation between samples, pinpoints candidate causal substitutions, and applies equally to the genomes of species and extinct lineages for which no functional data can ever be obtained.

Our work further illustrates a general strategy for studying gene regulatory evolution: learn regulatory grammar in the few species where functional genomics data are deep, then read it from the genomes of the many where only sequence is available. It can be extended along two axes: the cellular and developmental resolution of the functional data behind the model and the range of genomes it is applied to. Because human-specific regulatory changes act in particular cell types and developmental stages, fine-tuning the model on the growing number of single-cell RNA-seq and ATAC-seq datasets^24,55^, together with enhancer-to-gene maps at the same resolution^56^, would resolve changes that tissue-level profiles cannot capture, and the cell type and stage in which each acts. At the same time, genomes of modern and ancient humans, archaic hominins and other primates are becoming available at increasing scale^57–59^, so the same predictions can be made across many more individuals, species and lineages, complementing what population and evolutionary genetics infer from them. Together, these developments will extend the systematic investigation of gene regulatory evolution begun here to further cell types, developmental stages and lineages, bringing within reach a mechanistic understanding of how regulatory changes shaped the evolution of our species.

## Methods

### Human-lineage substitutions

All analyses are anchored on 16,958,523 biallelic single-nucleotide substitutions that are derived in the human reference genome (hg38) relative to the reconstructed human–chimpanzee ancestor (HCA). The ancestral reconstruction of Mangan et al., 2022^5^ was used as released. Briefly, ancestral states were estimated by maximum likelihood (gonomics: primateRecon) from a multiple alignment of the human, chimpanzee, bonobo, gorilla and orangutan genomes. The neutral rate of evolution was estimated from four-fold degenerate sites of the UCSC knownGenes set (PHAST msa_view), with branch lengths fit on a fixed-topology tree under a Jukes–Cantor model (PHAST phyloFit^60^). A base was considered present at the ancestral node when present in at least two species on two independent lineages connected to that node, and the posterior probabilities of A, C, G and T were then reconstructed. The most likely allele was accepted as ancestral only when its posterior probability was at least 99%; uncertain positions were set to N and excluded. The substitution set is therefore high-confidence rather than exhaustive.

### Cross-species bulk ATAC-seq, ChIP-seq and RNA-seq

#### Lymphoblastoid cell lines (LCLs)

Raw ATAC-seq and histone ChIP-seq data from LCLs of human, chimpanzee, gorilla, and orangutan^16^ were reprocessed with the ENCODE pipelines (atac-seq-pipeline v2.2.3; chip-seq-pipeline v2.2.2) using the official Docker images. Reads were aligned to each species’ own assembly: hg38 for human and the Ensembl toplevel assemblies Pan_tro_3.0, gorGor4 and Susie_PABv2 for the other great apes. ATAC reads were adapter-trimmed with TrimGalore^61^ (v0.6.10), aligned with Bowtie2^62^ (v2.3.4.3), separated from the mitochondrial contig, deduplicated with Picard^63^ (v2.20.7) and converted to Tn5-shifted tagAlign; peaks were called with MACS2^64^ (v2.2.4) per replicate, per pooled replicate and per pseudoreplicate, and reproducible peaks defined by irreproducible discovery rate (IDR). Histone ChIP reads were aligned with Bowtie2, filtered (-F 1804 -q 30) and deduplicated; fragment length was estimated by cross-correlation (phantompeakqualtools v1.2.1), peaks were called with MACS2 against each individual’s pooled input, and reproducible peaks defined by ENCODE naive overlap. Peaks were filtered against per-species uniquely mappable genome tracks and, for human, the ENCODE hg38 blacklist^65^.

##### Mappability tracks

Uniquely mappable genome tracks are available for hg38 but not for the non-human assemblies, so we generated them for every assembly to apply the same filter in each species. Single-read mappability was computed with Newmap (v0.2), a reimplementation of Umap^66^: an FM index was built over the genome, the shortest unique length computed at every position of the major chromosomes, and the result thresholded at the read length. Tracks were built at 50 bp (ATAC-seq) and 36 bp (histone ChIP-seq).

##### Epstein–Barr virus (EBV) decoy

Because LCLs are EBV-immortalized, the EBV genome (NC_007605) was added as an extra contig to every reference before alignment, so that EBV-derived reads map to it rather than to host sequence; these reads were then removed by the major-chromosome filter. The fraction of reads mapping to EBV was 0.16% (human), 0.20% (chimpanzee), 0.021% (gorilla) and 0.11% (orangutan).

#### Bulk RNA-seq across organs

Raw bulk RNA-seq data from cerebellum, heart, kidney, liver and testis of human, chimpanzee, bonobo and gorilla^30^ were reprocessed with the ENCODE RNA-seq pipeline (rna-seq-pipeline) on the official ENCODE image. Reads were aligned with STAR^67^ (v2.5.1b) to consensus gene annotations (below) and quantified with RSEM^68^ (v1.2.31) and kallisto^69^ (v0.44.0).

##### Consensus gene annotations

Because annotation quality differs between assemblies, and a gene with more annotated exons in one species accrues expression artifactually, expression was quantified against a consensus annotation projected across genomes, following Ma et al., 2022^70^. Union exon models of GENCODE protein-coding genes on human primary chromosomes were lifted reciprocally to each non-human assembly with UCSC liftOver (-minMatch = 0.1). Genes were retained when at least half of their exons mapped reciprocally and colinearly onto one chromosome (including unplaced scaffolds, so that assembly fragmentation does not discard genes), with per-exon length change below 50%, per-intron length change below five-fold and gene-body span change below four-fold. Surviving exons were bridged into a pre-mRNA gene-body model, so that gene identifiers and the gene set are shared across species. Analyses were restricted to 1:1 orthologs present in every species in a given contrast.

##### Normalization

Counts were converted to reads per kilobase million (RPKM) per species and placed on a common scale per tissue by anchor-gene median scaling, following the original study^30^. Within a tissue, orthologs with mean expression in the interquartile range were ranked within each sample, and the 1,000 genes with the lowest standard deviation of rank across samples were used as anchors, so that the scaling cannot be driven by genes that differ between species. Each sample was scaled so that its median expression over the anchors equaled the median across samples, and normalized RPKM was log_2_(RPKM + 1)-transformed. Genes were required to exceed a mean linear RPKM of 0.25, applied before transformation so that the transform cannot affect which genes pass.

### Cross-species single-nucleus RNA-seq (snRNA-seq) and ATAC-seq (snATAC-seq)

Published single-cell and single-nucleus data were used for cell-type-resolved expression and accessibility, processed per dataset as follows.

#### Liver

snRNA-seq and snATAC-seq data of liver from human, chimpanzee, bonobo, gorilla and orangutan were generated and processed by Yuan et al., 2025^18^. Of the six cell types recovered in all five species, B-cell lineage and hepatic stellate cells were excluded because their low cell numbers do not support differential accessibility testing, leaving hepatocytes, NK/T cells, macrophages, and endothelial cells. Pseudobulk gene counts were variance-stabilized per species with varistran^71^ (v1.1.0).

#### Cerebellum

snRNA-seq and snATAC-seq data of human cerebellum and single-nucleus multiome data of bonobo cerebellum were generated and processed by Sepp et al., 2024^41^ (human snRNA-seq) and Sarropoulos et al., 2026^11^ (all other data). Analyses were restricted to mature granule cells (GC_defined): the bonobo samples came from infant and toddler animals (6 months and 3 years), in which 84.9% of nuclei are granule cells against 1–4% for each other population, so no other cell type is represented well enough in both species. Per-sample pseudobulk counts were variance-stabilized per species with varistran and averaged per cell type, as for liver.

#### Cerebral organoids

Single-cell RNA-seq (scRNA-seq) and ATAC-seq (scATAC-seq) data of human and chimpanzee cerebral organoids were taken from Kanton et al., 2019^32^. For expression, the cross-species consensus count matrix and cell annotations were downloaded from ArrayExpress (E-MTAB-7552), cells annotated as radial glia at organoid stages were summed into pseudobulk profiles, and counts were variance-stabilized per species with varistran. For accessibility, per-cell alignments and annotations were downloaded from ArrayExpress (E-MTAB-8089 human, E-MTAB-8043 chimpanzee, E-MTAB-8083 bonobo); reads from neural progenitor cells were pooled per species, converted to coverage normalized by counts per million (CPM) in 200 bp bins, and lifted from hg19 (human) and panTro4 (chimpanzee and bonobo) to hg38 with UCSC liftOver.

#### Testis

snRNA-seq data of testis from human, chimpanzee, bonobo and gorilla were taken from Murat et al., 2023^54^, using the published count matrices, cell-type labels and germ-cell pseudotime. Per-cell expression was log_2_(1 + 10^4^ × count / nUMI), and germ cells were summarized in equal-width pseudotime bins.

### Differential chromatin accessibility between human and other great apes

Differential accessibility (DA) was analyzed in the bulk LCL ATAC-seq data and in each of the four liver cell types (snATAC-seq) with the same pipeline. Each pairwise comparison of human against one other great ape was tested separately: against chimpanzee, gorilla, and orangutan in LCLs, and against chimpanzee, bonobo, gorilla and orangutan in each liver cell type. Replicates were the two biological replicates per species in LCLs and the two to four individuals per species in liver.

Peak positions (narrowPeak summits for LCL IDR peaks; centers of the uniform 500 bp ArchR peaks for liver) were lifted reciprocally between hg38 and each non-human assembly with UCSC liftOver (-minMatch=0.1), retaining positions whose round trip returned to the same chromosome within 50 bp. Positions from both directions were merged in human coordinates (clustering within 250 bp), and each locus was represented by a 501 bp window in each genome, labeled by whether it derived from a human peak, a non-human peak or both. Analysis was restricted to autosomes, because replicates are not sex-matched between species. Reads were counted per window and replicate in each species’ own genome: Tn5-shifted tagAlign reads for LCLs, and for liver, primary alignments with MAPQ ≥ 30 reduced to 1 bp Tn5 insertion sites (5′ end shifted +4 bp on the plus and −5 bp on the minus strand using samtools^72^ (v1.20). GC content of the human window was computed with bedtools nuc^73^ (v2.30.0). Counts were filtered with edgeR::filterByExpr^74^ (v4.0.16) and normalized with cqn^75^ (v1.48.0), fitting a GC smooth with length as a fixed offset (lengthMethod=“fixed”), since windows are of near-constant length. The offsets, rescaled to a per-region geometric mean of one, were supplied to DESeq2^76^ (v1.42.1) as normalization factors, and log_2_ fold changes were shrunk with apeglm^77^ (v1.24.0). A window was classified as a human-specific accessibility gain at padj < 0.05 and log_2_FC > 1, a human-specific loss at padj < 0.05 and log_2_FC < −1, and conserved otherwise. Windows were retained only if the human window was ≥80% 50-mer uniquely mappable and outside the ENCODE hg38 blacklist^65^, and the non-human window was ≥80% uniquely mappable in its assembly (no blacklist exists for non-human assemblies). This filter was applied to the final orthologous windows rather than to peak sets, so that losses, which enter through non-human peak calls, are filtered by the same criterion.

Pairwise calls were then combined across species. Because the predictions concern substitutions that arose on the human lineage since the HCA, the quantity of interest is the change relative to that ancestor, which we inferred from the other great apes by parsimony: a change on the human lineage should differ from every other great ape, whereas a difference from a single species is more readily explained by a change on that species’ lineage or by noise. We therefore required differential accessibility against every species in a given set. This conservative choice trades sensitivity for a positive set unlikely to reflect changes on other lineages, and mirrors the sequence-side requirement that the ancestral allele be present in every species named. Because each comparison has its own set of testable loci, loci were matched across comparisons by overlap in human coordinates (bedtools multiinter^73^). A locus was called a human-specific gain against a set of species when called a gain in each of those comparisons, and a loss correspondingly. The conserved background for the same species set comprises loci called conserved in every comparison, after removing any locus called differentially accessible against any species in either direction. Each downstream analysis states the species set it uses.

### AlphaGenome substitution-effect prediction

We used the distilled AlphaGenome model^3^ (checkpoint alphagenome-jax-all_folds-v1), distilled from an ensemble of teacher models trained on all folds. Each substitution was scored with the recommended CenterMaskScorer (width = 501, aggregation_type = DIFF_LOG2_SUM) on a 1,048,576 bp (2^20^) input window centered on the variant, using the human output head. For each track, the raw score is the log_2_ of the predicted signal summed over the central 501 bp for the ancestral allele minus the same quantity for the human allele; a negative raw score therefore denotes a predicted human gain. Scores were recorded for all 167 human ATAC-seq and 305 human DNase-seq tracks from a single forward pass.

#### The AlphaGenome score

Raw DIFF_LOG2_SUM values are on track-specific scales and are not comparable between tracks or assays. Each raw score was therefore converted to an empirical percentile against AlphaGenome’s calibration set of common human variants (minor allele frequency > 0.01 in any gnomAD v3 population^57^), whose score distribution is provided with the model (calibration_scores.pb). The raw value was located in the track’s calibration quantile vector, giving AlphaGenome’s signed quantile q ∈ [−1, 1], which we report as the percentile p = 50(1 − q) ∈ [0, 100]. Because negative raw scores denote human gains, a strong predicted gain gives p near 100 and a strong predicted loss p near 0, and p = 50 means no predicted change. We refer to this percentile as the AlphaGenome score. A substitution was called a gain at the P-th percentile when p ≥ P (e.g., p ≥ 99 or p ≥ 99.9) and a loss when p ≤ 100 − P.

### Comparison with other sequence-based models

Four further models were scored on the same substitutions and evaluated identically: gkm-SVM and CNN classifier from Wang et al., 2026 in LCLs only, and Enformer and ChromBPNet in both datasets. All scores were oriented so that a high value denotes the direction being tested.

#### Enformer

Enformer^17^ was run through enformer-pytorch (v0.8.12) on a 196,608 bp input window centered on each substitution, reading all 674 human DNase-seq tracks from a single forward pass. As for AlphaGenome, the predicted signal was summed over the 128 bp output bins overlapping a 501 bp window centered on the substitution, and the effect was the log_2_ ratio of the ancestral to the derived sum. Evaluation used the matched-biosample track in LCLs and, in liver, a track selected per cell type as for AlphaGenome.

#### ChromBPNet

Bias-factorized ChromBPNet models^8^ (v1.0.1) were trained on human data: pooled, unshifted Tn5 tagAlign from LCLs, and separately the snATAC-seq data of each liver cell type. Training used called peaks and GC-matched negatives with five folds over the standard hg38 chromosome splits, as in the original study, with a bias model trained on fold 0 and reused across folds. Variant effects are the mean log_2_(ancestral/derived) over the five folds.

#### gkm-SVM

Gapped k-mer support vector machine classifiers were trained on human LCL peaks, one per fold, with default parameters (-t 2, l = 11, k = 7, d = 3, c = 1) using lsgkm^78,79^ (v0.1.2), with the same positives, GC-matched negatives and fold splits as ChromBPNet. Inputs were 1,000 bp windows, centered on the peak summit for training and on the substitution for scoring. The variant effect is the mean over folds of score(derived) − score(ancestral).

#### CNN classifier from Wang et al., 2026

Variant effects on LCL chromatin accessibility were scored with the published convolutional neural network (CNN) classifier of Wang et al., 2026^13^. The model architecture (six one-dimensional convolutional layers in three convolution–convolution–max-pool blocks, adaptive average pooling and two fully connected layers with a sigmoid output) and the one-hot encoding were reimplemented following the authors’ code, and their released weights for the four human LCL open-chromatin replicate models were loaded unchanged. Each model was applied to 1,000 bp windows centered on each substitution, and the variant effect is the difference between the sigmoid outputs for the derived and ancestral sequences, averaged over the four replicate models.

### Benchmarking against measured differentially accessible (DA) peaks

Performance was evaluated per DA-tested region. Positives are regions with a human-specific gain (or loss) against every species in the given set, and negatives the conserved background for the same set. Each region was assigned the maximum direction-oriented variant score over its substitutions, reflecting the expectation that a single substitution can largely account for a change; an additive aggregation gave equivalent performance. Regions without a scored substitution were dropped. Precision–recall curves, average precision and the prevalence baseline were computed with scikit-learn^80^ (v1.6.1), and comparisons were restricted to conditions fully covered by every model. In LCLs, precision reached 0.19 at an AlphaGenome score threshold of 99 and 0.65 at 99.9, 7- and 25-fold higher than expected by chance, at 57% and 26% recall, respectively.

Requiring differential accessibility against every species is harder to satisfy for losses than for gains, because a loss requires the region to be reliably detected as accessible in each non-human genome, each with its own sequencing depth and mappability, whereas a gain requires detection in human only. Fewer losses therefore pass, and the largest species set retaining a usable positive set differs by direction. In LCLs, gains were called against chimpanzee, gorilla, and orangutan, and losses against chimpanzee and gorilla; in liver, gains against chimpanzee, bonobo, gorilla and orangutan, and losses against chimpanzee, bonobo and gorilla.

#### Track selection for AlphaGenome and Enformer

Both models output prediction on many tracks. For the LCL benchmark, the matched track (GM12878) was used for each model, without reference to performance. For the liver benchmark, each cell type used the track with the highest mean auPR across the species comparisons (Supplementary Table 2). Because this selection was made on the evaluation data, it was optimistically biased. As a sensitivity analysis, selecting the ontology-matched track without reference to performance still left AlphaGenome above ChromBPNet in the same seven of the eight cell type × condition comparisons (Supplementary Table 2). For Enformer, whose tracks are donor-specific and match two to five tracks per cell type, the same procedure reduced its advantage over ChromBPNet from six of eight comparisons to three, so its margin depended on how tracks were selected.

### Cross-species accessibility and histone profiles at predicted loci

For a given AlphaGenome score threshold, predicted sites were substitutions passing the threshold that lie in a human peak (gains) or in a peak of every required non-human species (losses), carry the ancestral allele in every required species, and pass the window quality filter used for DA analysis (±250 bp). These sites are drawn genome-wide rather than from DA-tested peaks. Only the species used to ascertain a site set are shown.

Each site was lifted to each assembly, and that species’ coverage was read in a ±500 bp (accessibility) or ±2,500 bp (histone modifications) window around the orthologous position; only loci measurable in every species shown were plotted. To make tracks comparable across species, CPM-normalized coverage was rescaled by the signal of constitutive peaks, as in previous cross-cell-type analyses^81,82^: each non-human species received one multiplicative factor, chosen so that strong, shared, non-differential windows (peaks in both genomes, |log_2_FC| < 0.5, top 1% by base mean) have the same mean height as in human. Loci are ordered by human mean signal, with one color scale shared across species. For histone panels, sites were re-filtered against the 36-mer mappability track and classified with human ChIP-seq peaks as promoter-like (H3K4me3 and H3K27ac) or enhancer-like (H3K27ac and H3K4me1). Matched background sites are substitutions in the same peak sets whose predicted effect is below the 1st percentile in both directions (AlphaGenome score under 1 for gained and for lost).

In LCLs, 1,044 regions contained at least one substitution with an AlphaGenome score ≥ 99.9; 82.7% of them were more accessible in human than in each of chimpanzee, gorilla, and orangutan, against 31.1% of matched background regions (Fig. 1d; Extended Data Fig. 1c; OR = 10.6; *P* = 2 × 10^−76^, Fisher’s exact test). Likewise, 95.6% of 508 regions with predicted losses were less accessible in human than in both chimpanzee and gorilla, against 40.1% of matched background regions (Extended Data Fig. 1c; OR = 33.8; *P* = 5 × 10^−74^, Fisher’s exact test). In liver cell types, 60.8–79.5% of predicted gains and 62.5–92.0% of predicted losses were confirmed by measured accessibility (Extended Data Fig. 1d).

### Genome-wide prediction of human-specific cis-regulatory gains

Human-specific cis-regulatory gains were predicted across all 305 human DNase-seq tracks, comprising 95 tissues from 33 organs (60 adult and 35 developmental), 103 primary or in vitro-derived cell types and 107 cell lines. A substitution was called a gain in a tissue when its AlphaGenome score in that track passed the threshold and it lay within open chromatin of the same biosample. This requirement applies to gains only, since a human gain should coincide with a human peak whereas a loss is not expected to. Regions were defined by merging substitutions with gains within 501 bp. With a threshold of 99.9, this yields 146,513 substitutions in 128,646 regions and 1,020,307 tissue × region calls across the 305 tissues.

#### Open chromatin

The 305 DNase-seq tracks are backed by 535 ENCODE experiments^19^. For each experiment, one GRCh38 narrowPeak file was taken from the ENCODE portal, preferring the released, pooled, reproducible call flagged as the experiment’s default, and otherwise the highest-priority reproducible output. For each track, peak files were combined with bedtools multiinter^73^ (v2.30.0), and intervals supported by at least min(2, n) of the track’s n experiments were retained; 238 of the 305 tracks are backed by a single experiment and thus use its reproducible call. This definition of open chromatin is used throughout.

### Tissue programs

For clustering, a region × track matrix was built in which each region took, in each track, the AlphaGenome score of its strongest substitution among those in that track’s open chromatin, transformed to a signed log scale, −log_10_(2 − P/50) for gains and log_10_(2 − P/50) for losses, where P is the AlphaGenome score. The scale is antisymmetric, so gains and losses of equal score take equal and opposite values and a score of 50 maps to zero. Entries where the region was not in the track’s open chromatin were set to zero (no predicted change) for clustering and are shown in a distinct background color.

Tracks were clustered hierarchically (average linkage, correlation distance), and the dendrogram was cut into K groups. Regions were not clustered independently: each was assigned to the track cluster in which its predicted gain was strongest, or to a broad class when gained in at least 10% of tracks. To choose K, two independent criteria were evaluated over K ≤ 100, each bootstrapped over regions: the silhouette score, which measures how well the partition matches the structure of the data without external information, and the adjusted mutual information with ENCODE’s organ annotation of the biosamples, corrected for chance agreement. Because differences between neighboring K are often smaller than the bootstrap error, we applied the one-standard-error rule to each, taking the smallest K within one standard error of the best. Both criteria returned K = 50 for all 305 tracks.

Clustering all 305 tracks yielded one large cluster combining cell lines and primary cells irrespective of their tissue of origin, except immune primary cells. Because this grouping likely reflects culture state rather than tissue identity, we removed all cell lines and all non-immune primary cells. The remaining 144 tracks comprise 95 primary tissues, 19 in vitro-differentiated cell types and 30 immune primary cell types, for which both criteria returned K = 25, yielding 25 tissue programs plus the broad class. All subsequent analyses use this 144-track panel (the 305-track clustering is shown in Extended Data Fig. 2). With a threshold of 99.9, the 144-track panel contains 79,135 substitutions with gains in 71,345 regions.

### Transcription factor motif enrichment

Motif instances were taken from ENCODE-GRAMMAR (Gene Regulatory Atlas of sequence Models, Motifs, Annotations, and Rules)^20^ for the experiments underlying each track. These annotations were produced with ChromBPNet^8^, DeepSHAP^83^, TF-MoDISco^84^ and FiNeMo^85^, with one model per experiment, and were downloaded as bigBed files. Instances were summarized per track as for open chromatin, retaining those supported by at least min(2, n) experiments. Motif calls are available for 124 of the 144 tracks, to which the analysis was restricted.

For each program, overlapping instances of the same motif across the program’s tracks were merged, so that each site is counted once. The foreground comprised merged instances containing at least one substitution with a gain in that program, and the background comprised instances containing at least one substitution from a cluster-balanced pool but none with a gain. The pool draws equal numbers of substitutions in open chromatin from each of the 25 programs (as many as the smallest program allows), so that differences in assay depth between programs do not set the background, and includes all substitutions with gains, so that each foreground is contained in the background. Conditioning the background on carrying a substitution makes the test about gains rather than about how much human-lineage divergence a motif family’s sites carry, and removes the bias from longer motifs catching more substitutions. Motif labels were collapsed into families by removing the confidence suffix, keeping activating and repressive directions separate. Enrichment is reported as odds ratios with *P* values from Fisher’s exact test.

### Functional enrichment of nearby genes

Each substitution in the pool was extended to a 501 bp interval, and intervals were merged once over the whole pool to define a common region universe, so that each program’s foreground is an exact subset of the background and region length does not differ systematically between programs. For each program, GREAT (Genomic Regions Enrichment of Annotations Tool)^86^ was run on the regions with its gains against this background using rGREAT^87^ (v2.0.2), and terms were considered significant at padj < 0.05 (gene-based hypergeometric test).

### Classification of human-specific cis-regulatory gains by mode of gain

For each region with a predicted gain assigned to one of the 25 programs, we asked whether it is also accessible in human in other programs and, if so, whether its substitution is predicted to increase accessibility there as well. A region was considered accessible in a program if its substitution lay within an ENCODE DNase-seq peak in at least one experiment of at least one of the program’s tracks. In each such program, we took the highest AlphaGenome score among the tracks in which the region is accessible, and counted the program as gained when this score was ≥99.9. The program in which the gain was called was excluded. Regions accessible in no other program, or only in other programs that were all gained, were classified as de novo gains; regions accessible in at least one other program without a comparable predicted gain were classified as co-option, with accessibility in that program inferred to predate the substitution.

Motif instances were taken from ENCODE-GRAMMAR^20^ as above, using the cross-sample motif vocabulary, pooled over each program’s tracks, and restricted to instances with a positive contribution to accessibility. CTCF instances were excluded given their enrichment in the broad class, and families were compared at the level of the base motif name. Of the 25 programs, 22 had motif calls, covering 67,303 of the 68,031 classified regions. For each region, we recorded the motif families with an instance covering the substitution in the program in which the gain was called, and those with an instance elsewhere in the 501 bp region. De novo gains were compared using the presence of each family at the substitution (Extended Data Fig. 3a), and co-opted regions using its presence elsewhere in the region (Extended Data Fig. 3b), each against de novo gains confined to one program. Within co-opted regions (Extended Data Fig. 3c), each family’s share of instances at the substitution was compared with its share elsewhere, using instance counts so that the larger number of instances elsewhere does not shift all families in the same direction. In all three comparisons, odds ratios were pooled over programs by the Mantel–Haenszel method with Robins–Breslow–Greenland confidence intervals, and *P* values were adjusted by the Benjamini–Hochberg method within each panel.

### Properties of predicted accessibility-gained CREs

Features were computed for every open-chromatin peak of every track: GC fraction; CpG observed/expected, (n_CG × width) / (n_C × n_G); log_10_ distance from the peak midpoint to the nearest protein-coding transcription start site (TSS); mean constraint across 470 mammals (phastCons^88^ 470-way); breadth of accessibility (mean fraction of tracks whose peaks cover each base); number of HCA substitutions; log_10_ peak width; and within-track percentile of peak signal. Peaks were required to have constraint scores over at least half their length.

For each tissue, a logistic regression was fit over the peaks of its open chromatin, with the outcome being whether a peak contains at least one substitution with a gain in that tissue. Because peaks with more substitutions have more chances to contain a gain, the log_10_ number of substitutions entered as an exposure term, so that other coefficients are interpreted at fixed mutational opportunity. Accessibility breadth, TSS distance, CpG observed/expected and constraint were the features of interest; GC fraction, signal percentile and peak width were included as covariates. Continuous terms were z-scored within track. No variance inflation factor exceeded 5. Tracks with fewer than 50 gains were not fit (chondrocyte, CL:0000138, 27 substitutions in 26 regions; mesenteric fat pad, UBERON:0015143, 48 regions after coverage and autosome filters), leaving 142 tracks.

The 142 per-tissue coefficients were combined by random-effects meta-analysis, a standard approximation to a generalized linear mixed model^89^, which would be infeasible here (eight random slopes over ∼20 million rows); agreement was checked by fitting the mixed model on a stratified subsample. The pooled effect μ is the mean of tissue coefficients weighted by 1/(SE^2^ + τ^2^), with between-tissue variance τ^2^ estimated by the Paule–Mandel method^90^, so that precise tissues count more only until genuine between-tissue disagreement dominates. Because tissues share peaks and substitutions and are therefore not independent, the standard error of μ was estimated by a leave-one-cluster-out jackknife over the 25 programs. Effects are reported as odds ratios, exp(μ), with 95% intervals exp(μ ± 1.96 × SE_jackknife).

To show each feature in its own units, regions with gains were compared with a background reweighted to match them on all other covariates by inverse probability weighting^91^: within each tissue, a main-effects logistic propensity model estimated the probability of a gain given all covariates except the focal one, and regions without gains were weighted by w = p/(1 − p), which makes the weighted background’s covariate distribution equal that of regions with gains^92^. This retains all regions with gains and achieved a median absolute standardized mean difference of 0.009.

#### Comparison with ENCODE candidate CRE (cCRE) annotation

Each region with a gain in the 144-track panel was assigned the class of the ENCODE cCRE (v4, GRCh38)^93^ containing its representative substitution, or labeled unannotated. The background comprised merged DNase-seq peaks of the 144 tracks carrying at least one human-lineage substitution outside regions with gains, each represented by one randomly drawn substitution and reweighted to match regions with gains on peak strength (within-experiment signal percentile of the strongest covering peak, in 5-percentile bins) and peak width (eight bins). Enrichment per class was the odds ratio against all other classes, with Woolf 95% confidence intervals and Fisher’s exact test.

#### Comparison with HARs and HAQERs

Human accelerated regions (HARs)^4,26^ and human ancestor quickly evolved regions (HAQERs)^5^ were intersected with the predicted gained element set in hg38 coordinates.

### Proximity analysis of gained elements

Clustering was assessed as a one-dimensional marked point process, in which points are candidate regions at fixed genomic positions and the mark is a gain. For each distance bin, the statistic is the number of pairs at that distance with both ends marked, divided by the mean of the same count over permutations of the marks, so that a value of one indicates spatial independence. Marks were permuted within non-overlapping 1 Mb blocks, preserving the number of gains per megabase and thus regional structure (e.g., gene deserts, isochores, enhancer-dense domains, replication timing and coverage) without enumerating covariates. Chromosome-wide permutation, 100 kb blocks and covariate quartile strata were computed as sensitivity checks, and block sizes from 500 kb to chromosome-wide are shown in Extended Data Fig. 6a. Nulls in which fewer than 60% of marked regions share a permutation cell with an unmarked region cannot move those marks and were dropped. Distance bins start at 3 kb, because substitutions within 501 bp are merged by construction and the median gap between neighboring regions is 0.9 kb, which deflates pair counts at shorter distances. Each null is informative only at separations well below its block length.

For measured differential accessibility, gains and losses were placed on one point set and a three-level label (neither, gain, loss) was permuted jointly, preserving both marginal counts so that gain–gain, loss–loss and gain–loss pairs share one null. This distinguishes coordinated directional change from correlated noise, since DA errors are spatially correlated by construction (neighboring peaks share fragments, coverage, mappability and copy number) but blind to direction, and would raise all three terms together. Measured gains were those called against all four other great apes in liver and against chimpanzee, gorilla, and orangutan in LCLs; losses against chimpanzee, bonobo and gorilla in liver and against chimpanzee and gorilla in LCLs, since the deepest species set leaves too few losses. Cell types were pooled: pairs were counted within each cell type, and observed and permuted counts summed across cell types per permutation, which also defines the confidence intervals. For predicted gains, the background was restricted to each program’s own accessible regions, matching the measured analysis, in which the background is each cell type’s own conserved peaks. To test whether the weaker clustering of predicted gains is explained by prediction error alone, we simulated predicted sets with the model’s set size and number of correct calls, drawing hits at random from measured gains and misses at random from the remaining peaks, so that errors are spatially unbiased, and compared the observed pair count with 10,000 simulations.

Across LCLs and the four liver cell types, measured gain–gain and loss–loss pairs were enriched at 3–10 kb (pair enrichment 1.91 and 1.85, respectively; *P* < 1 × 10⁻⁴), whereas gain–loss pairs were strongly depleted (pair enrichment 0.18; Extended Data Fig. 6a). Predicted gains also clustered, although more weakly (pair enrichment 1.11 at 3–10 kb; *P* < 1 × 10⁻⁴). The predicted set yielded 25 close pairs, compared with 36 expected from the simulations (one-sided *P* = 0.038; Extended Data Fig. 6b), indicating that the weaker clustering exceeds what prediction error alone would explain.

#### Recovery of measured gains

For measured gains in LCLs and liver cell types, we asked how often AlphaGenome recovers each as a function of the number of other measured gains within 30 kb, the range over which pair enrichment is detectable (present at 3–10 and 10–30 kb, absent at 30–100 kb). The predicted set was size-matched to the measured one per cell type. Because cell types differ in baseline recovery, plotted values were standardized by fitting a logistic regression on neighbor count and cell type and averaging predictions over the observed cell-type mix, with bootstrap confidence intervals resampled within cell type. The trend was tested by entering neighbor count as a linear score in a logistic regression with cell-type fixed effects. Recovery decreased from 45.0% for isolated gains to 28.5% for those with two or more neighboring gains (OR = 0.68 per neighbor; *P* = 1.0 × 10^−6^, Wald test; Extended Data Fig. 6c).

#### Sequence support within clusters

Clusters were defined as groups of at least two measured gains whose midpoints lie within 20 kb. Within clusters containing both recovered and unrecovered gains, the accessibility change of recovered gains was compared with that of their unrecovered neighbors by a Wilcoxon signed-rank test (*P* = 3.7 × 10^−9^), and the two classes were compared by Fisher’s exact test for carrying a human-lineage substitution within a motif instance, per cell type and pooled. Motif instances came from each cell type’s own biosample for hepatocytes, LCLs and endothelial cells; the biosamples for NK/T cells and macrophages lack motif calls, so the next-best biosamples in the track selection were used (effector memory CD8-positive alpha-beta T cell and CD14-positive monocyte, at a cost of 0.8% and 4.0% of mean area under the precision–recall curve), and the macrophage estimate is not interpreted alone. Whether clusters contain a recovered gain more often than expected was tested by permuting recovery labels among each cell type’s clustered gains (preserving the number recovered per cell type; 10,000 permutations, one-sided).

### Linking predicted gained CREs to genes

Regions with predicted gains were linked to genes with ENCODE-rE2G^29^. For each dataset, the human CREs, rE2G maps and AlphaGenome predictions were matched to the tissue or cell type of the expression data as closely as available human data allowed (Supplementary Table 7). For the 144-track panel, the map for each track is its rE2G prediction averaged over experiments. Because one merged region can span several rE2G elements, each region–gene pair took the highest link score among them, and a region was considered linked to a gene if it had any rE2G link to it. For each gene, the gained rE2G burden is the sum of link scores over its linked regions with gains, and the corresponding sum over its other linked CREs was computed as a measure of remaining regulatory input. Because the burden aggregates over many regions, gains were called with an AlphaGenome score threshold of 99 for these analyses, favoring sensitivity over precision.

To compare the burden with expression differences, we used cell-type-resolved datasets from liver^18^ and cerebellum^11^, which have matched accessibility data, and bulk RNA-seq from cerebellum, heart, kidney, liver and testis^30^, for which ENCODE accessibility data from the matched tissue were used. For single-cell datasets, the CRE universe is the cell type’s own accessibility data, and predictions and links come from the closest ENCODE biosample. For liver, the CRE universe is each cell type’s merged snATAC-seq peak set, with predictions from the best-matching DNase-seq track (right lobe of liver for hepatocytes, immature natural killer cell for NK/T cells, suppressor macrophage for macrophages, and dermis blood vessel endothelial cell for endothelial cells) and links from the same biosample, except for hepatocytes, for which the left lobe of liver map was used. For cerebellum, the CRE universe is the merged reproducible snATAC-seq peak set of mature granule cells, with predictions from the cerebellar cortex track and links from the whole-cerebellum map. For tissue-level datasets, the DNase-seq biosample was selected by testing each relevant biosample with the same link set and ranking by Spearman correlation. Genes were binned by gained rE2G burden (deciles for single-cell and quintiles for tissue-level datasets, since bulk contrasts carry more measurement error). Single-cell bins are shown as notched boxes (interquartile range; whiskers, 10th–90th percentiles; notch, 95% confidence interval of the median), and tissue-level bins as the median expression difference with a percentile bootstrap confidence interval (2,000 resamples). Expression differences were median-centered within each dataset. A gene was called human-upregulated when its centered difference was above zero, so that the comparison is with the typical gene of the dataset rather than with equal expression between human and other great apes. The fractions of human-upregulated genes in the top and bottom burden bins were compared with a two-sided Fisher’s exact test.

To test whether other candidate sets of human-lineage regulatory regions show the same relationship with expression, we repeated the analysis with CREs overlapping each of four published sets: 2,698 HARs^4,26^, 1,581 HAQERs^5^, 13,027 human predicted increased chromatin accessibility regions (hPICAs; union across all 111 cell types)^13^, and 7,367 hominin-lineage-gained CREs^14^. A CRE was counted when overlapping at least one region of a set, and the burden was summed over these CREs instead of regions with gains, with all other inputs unchanged. Regions with predicted gains, processed identically as a reference, reproduced the results above. The burden was entered in the same regression model without standardization, because the sets differ greatly in sparsity.

#### Properties of genes with a high gained rE2G burden

Gene properties were taken from external resources independent of the expression data. Tissue specificity (tau^94^) was computed on log_2_(nTPM + 1) across the 51 tissues of the Human Protein Atlas consensus dataset^95^. Dosage sensitivity was s_het from Zeng et al., 2024^96^, encoded as membership in the top decile, since the association with burden was confined to the most constrained genes. Subcellular localization was encoded as three indicators from Gene Ontology cellular-component annotations^97,98^: nucleus (GO:0005634), extracellular (GO:0005615 or GO:0031012) and integral membrane (GO:0016021 or GO:0005887), with unannotated genes coded as zero. Genes without an s_het estimate were excluded. Within each track, genes with at least one rE2G link and complete annotation were ranked by burden, and genes in the top decile were compared with the rest by logistic regression on the four properties and regulatory complexity (continuous terms z-scored within track). Tracks with fewer than 50 regions with gains or fewer than 500 genes were excluded. Coefficients were pooled by random-effects meta-analysis with leave-one-cluster-out jackknife intervals, as above, and consistency was summarized as the fraction of tissues expected to share the direction of each effect. To show each property in its own units, genes outside the top decile were reweighted by propensity to match the top decile on all other terms, as above, and cumulative distributions were averaged over tracks.

### Candidate loci

For each candidate gene, the plotted window spans the gene body and its linked regions with gains carrying a reproducible rE2G link, padded by a tenth of that span; for datasets with only the full link set, where any-link anchoring would span megabases, the window was based on the gene’s few highest-scoring regions instead. Arcs show the links used to define the window, plus full-set links of any region with a gain lacking a reproducible link, so that every region marked as linked has an arc.

Cross-species accessibility was displayed on hg38 by tiling the window, lifting each tile to each species, reading that species’ coverage and writing it back on the hg38 axis. Tiles that failed to lift, or lifted to more than three times their length, were left missing rather than set to zero, since absent alignment is not absent accessibility, and plotting it as zero would create apparent human gains where alignment is worst. For the organoid radial glia panels, all three species were lifted once at the fragment level to hg38 (human from hg19; chimpanzee and bonobo from panTro4). Each track was normalized to its own library size.

#### Per-base contribution to accessibility

Per-base contribution scores were computed by in silico saturation mutagenesis: every base within 50 bp of a substitution with a gain was mutated to each alternative, and its importance was the negative mean log_2_ ratio of mutated to reference predicted accessibility, so that upward letters mark bases the human sequence requires for accessibility and downward letters bases whose mutation would increase it. Motif instances created by substitutions were annotated manually, by comparing the high-contribution pattern surrounding each substitution against known TF binding profiles using Tomtom^99^, and against the motif annotations of the same region in the AlphaGenome Atlas^9^.

#### In silico ancestralization

For per-region predictions, the region’s human alleles were reverted to their reconstructed ancestral states and AlphaGenome was re-run on the dataset’s accessibility track, with a ∼1 Mb input window centered on the region and the signal summed over the region. For gene-level predictions, the substitutions in all linked regions with gains were reverted simultaneously in a single run, and the output read on the matching RNA-seq track.

### Identifying modern-human-specific changes using archaic genomes

Substitutions were dated with seven high-coverage archaic genomes: five Neandertals (Altai^42^, Chagyrskaya Phalanx^43^, Vindija 33.19^44^, Goyet Q56-1 (European Nucleotide Archive PRJEB98484) and Denisova 17^45^) and two Denisovans (Denisova 3^46^ and Denisova 25^47^). Genotypes were read from per-individual consensus sequences generated without reference bias, in which unsequenced bases are encoded as N, so that uncallable sites are distinguished from called ones. This matters because 35.1% of HCA substitution sites cannot be genotyped in any archaic genome, largely owing to their short reads and lower quality.

A substitution was classified as modern-human-specific when every genotyped archaic genome carried the ancestral allele, with at least one Neandertal and one Denisovan genotyped; as shared with archaic hominins when at least one genotyped archaic carried the derived allele and none the ancestral; as mixed when genotyped archaics disagreed; and as unresolved when no archaic could be genotyped. A region was modern-human-specific when any of its substitutions with gains was, and shared with archaic hominins when none was modern-human-specific and at least one was shared. Analyses were restricted to autosomes, as the archaic genotypes are autosome-only.

Because missing data differ between sets (substitutions with gains are better covered than the genome average, 18.6% vs. 35.1% uncallable, because open chromatin is more mappable), comparisons between sets use the fraction modern-human-specific among modern-human-specific plus shared, rather than raw percentages, which would conflate differences in coverage with differences in age. Modern-human-specific does not imply fixed: genome-wide, 95% of modern-human-specific substitutions with gains are still segregating in African populations of the 1000 Genomes Project. Derived allele frequencies in the five African populations are therefore reported alongside dating, and the fixed or nearly fixed subset was analyzed separately.

The proportion of modern-human-specific regions relative to those shared with archaic hominins did not differ significantly between the 25 tissue programs and the broad class (*P* = 0.11, chi-squared test; all per-program FDR q ≥ 0.27), suggesting that no tissue program experienced a marked change in the rate of cis-regulatory gains before versus after the split between modern humans and archaic hominins.

### GC-biased gene conversion

GC-biased gene conversion (gBGC) preferentially transmits the strong (G or C) allele at weak/strong heterozygous sites, accelerating the fixation of weak-to-strong substitutions, and can therefore make a set appear both older and at higher frequency than neutral expectation^100^. Each autosomal human-lineage substitution was assigned a strand-collapsed mutation class (weak-to-strong, strong-to-weak at a CpG, strong-to-weak elsewhere, and the gBGC-insensitive A↔T and G↔C), with CpG status taken from the hg38 flanking base. Contrasts were repeated within mutation class and combined by the Mantel–Haenszel estimator, and separately restricted to the two gBGC-insensitive classes.

gBGC tracts were called with phastBias^101^ on a six-ape alignment (human, chimpanzee, bonobo, gorilla, orangutan, gibbon) extracted from the Zoonomia Cactus alignment^102^ with hal2maf (-- onlyOrthologs), against a REV neutral model fit to the same species and with human as foreground (B = 3, expected tract length 1 kb, ρ = 0.31, conserved element length 45, conserved target coverage 0.3). The autosomes were scanned as 2,705 uniform 1 Mb tiles (2.7 Gb). The gBGC target coverage was estimated by expectation maximization on the 400 largest tiles (median 0.81%) and then held fixed, as per-tile estimates are unstable and a tile estimated high calls disproportionately many tracts. This gave 11,132 tracts over 0.40% of the scanned sequence, holding 146,629 of 16,221,343 substitutions on autosomes, which were excluded in the corresponding rows.

Two-by-two contrasts used Fisher’s exact test, stratified contrasts the Mantel–Haenszel odds ratio with Robins–Breslow–Greenland intervals, and substitution bias and tract fractions 95% percentile bootstrap intervals from 1,000 resamples of blocks (gained region, HAR or HAQER, or 1 Mb tile). Substitutions with predicted gains carried more gBGC than the genome average (f_W→S_ 0.499 versus 0.463, 1.27% versus 0.90% inside a tract) but far less than HARs (0.586, 6.9%) or HAQERs (0.524, 15.6%), and neither dating result changed under any correction, holding at an odds ratio of 0.80 for the depletion of modern-human-specific substitutions and 1.40 for the excess of high-frequency derived alleles (Extended Data Fig. 12).

### Ancient selective sweeps

To test whether predicted cis-regulatory gains fall preferentially in regions that experienced ancient positive selection, we used the candidate sweep regions of Peyrégne et al., 2017^51^, a scan that targets fixation events on the modern human lineage after the split from Neandertals and Denisovans but before the differentiation of modern human populations. The method identifies regions where both archaic lineages fall outside present-day human variation over a stretch longer than expected under neutral lineage sorting, indicating fixation too rapid for recombination to break up the surrounding haplotype. We used the lenient set of 314 regions (supported by at least one of the two recombination maps), lifted from hg19 to hg38 with liftOver, all mapping without fragmentation. Because these regions are ascertained in part from fixed derived differences between modern humans and archaics, the comparison used as background fixed modern-human-specific substitutions that were not gained and lay in the 144-track open chromatin union, matching both age and chromatin context. Significance was assessed by Fisher’s exact test.

## Acknowledgements

We thank Duncan Odom, Oliver Stegle, Ralph Grand, Alexander Sasse, Mari Sepp, Manu Saraswat, Jiawei Luo, Nils Trost, Marwan Elkrewi, Anamaria Elek, and all other members of the Kaessmann lab for discussions and Kathrin Hall, Julia Schmidt and Céline Schneider for assistance. The computational cluster bwForCluster Helix of Heidelberg University Computational Center is supported by the state of Baden-Württemberg through bwHPC and the German Research Foundation (INST 35/1597-1 FUGG). The authors gratefully acknowledge the data storage service SDS@hd supported by the Ministry of Science, Research and the Arts Baden-Württemberg (MWK) and the German Research Foundation (DFG) through grant INST 35/1803-1 FUGG and INST 35/1804-1 LAGG. T.Y. was supported by the Takenaka Scholarship Foundation. This study was supported by grants from the European Research Council (grant agreement nos. 101019268) and the NOMIS Foundation (no. 2024-A-0035) to H.K.

## Author contributions

T.Y., I.S., and H.K. conceived the study. T.Y. analyzed data with support from X.Y. H.Z., V.S., M.L.A., and S.P. provided archaic genome data. H.K. supervised the study. H.K. provided funding. T.Y. and H.K. wrote the manuscript, with critical review by I.S., V.S., and S.P. All authors approved its final version.

## Data availability

### Data generated in this study

Predicted human-gained cis-regulatory elements, per-substitution AlphaGenome scores and the source data underlying each figure are available and interactively explorable at https://apps.kaessmannlab.org/human_specific_cre.

### Previously published sequencing data

Bulk RNA-seq of human and great-ape tissues (cerebellum, heart, kidney, liver, and testis) from Brawand et al., 2011^30^ is available from the Sequence Read Archive (SRA) under accession SRP007412. LCL ATAC-seq and ChIP-seq from García-Pérez et al., 2021^16^ is available from the SRA under accession SRP220230. Cerebral organoid data from Kanton et al., 2019^32^ are available from ArrayExpress, comprising scRNA-seq of human and chimpanzee organoids (E-MTAB-7552) and scATAC-seq of human (E-MTAB-8089), chimpanzee (E-MTAB-8043) and bonobo (E-MTAB-8083) organoids. Testis snRNA-seq from Murat et al., 2023^54^ is available from ArrayExpress for human (E-MTAB-11063), chimpanzee (E-MTAB-11064), bonobo (E-MTAB-11067) and gorilla (E-MTAB-11065). Liver data from Yuan et al., 2025^18^ are available from ArrayExpress, comprising snRNA-seq of human (E-MTAB-15545), chimpanzee and bonobo (E-MTAB-15548) and gorilla and orangutan (E-MTAB-15549), and snATAC-seq of human (E-MTAB-15541) and chimpanzee, bonobo, gorilla and orangutan (E-MTAB-15546). Cerebellar data comprise human snRNA-seq from Sepp et al., 2024^41^, available from heiData under accession QDOC4E, and human snATAC-seq together with bonobo single-nucleus multiome from Sarropoulos et al., 2026^11^, available from heiData under accession GDSZG9.

### ENCODE resources

Open-chromatin peak calls and signal tracks, CRE to gene links from ENCODE-rE2G^29^ and TF binding motif instance annotations from ENCODE-GRAMMAR^20^ were obtained from the ENCODE portal (https://www.encodeproject.org).

### Archaic hominin genomes

High-coverage Neandertal genomes are available at https://ftp.eva.mpg.de/neandertal/ and Denisovan genomes at https://ftp.eva.mpg.de/denisova/, with the exception of Goyet Q56-1, available from the European Nucleotide Archive (ENA) under accession PRJEB98484.

### Model weights

AlphaGenome model weights^3^ are available from Hugging Face (https://huggingface.co/collections/google/alphagenome). Enformer model weights^17^ were obtained through enformer-pytorch v0.8.12. The CNN classifier from Wang et al., 2026^13^ is available at https://doi.org/10.5281/zenodo.18925037.

## Code availability

All original code is available at https://gitlab.com/kaessmannlab/human-specific-cres.

## Extended Data Figures

**Extended Data Fig. 1:**
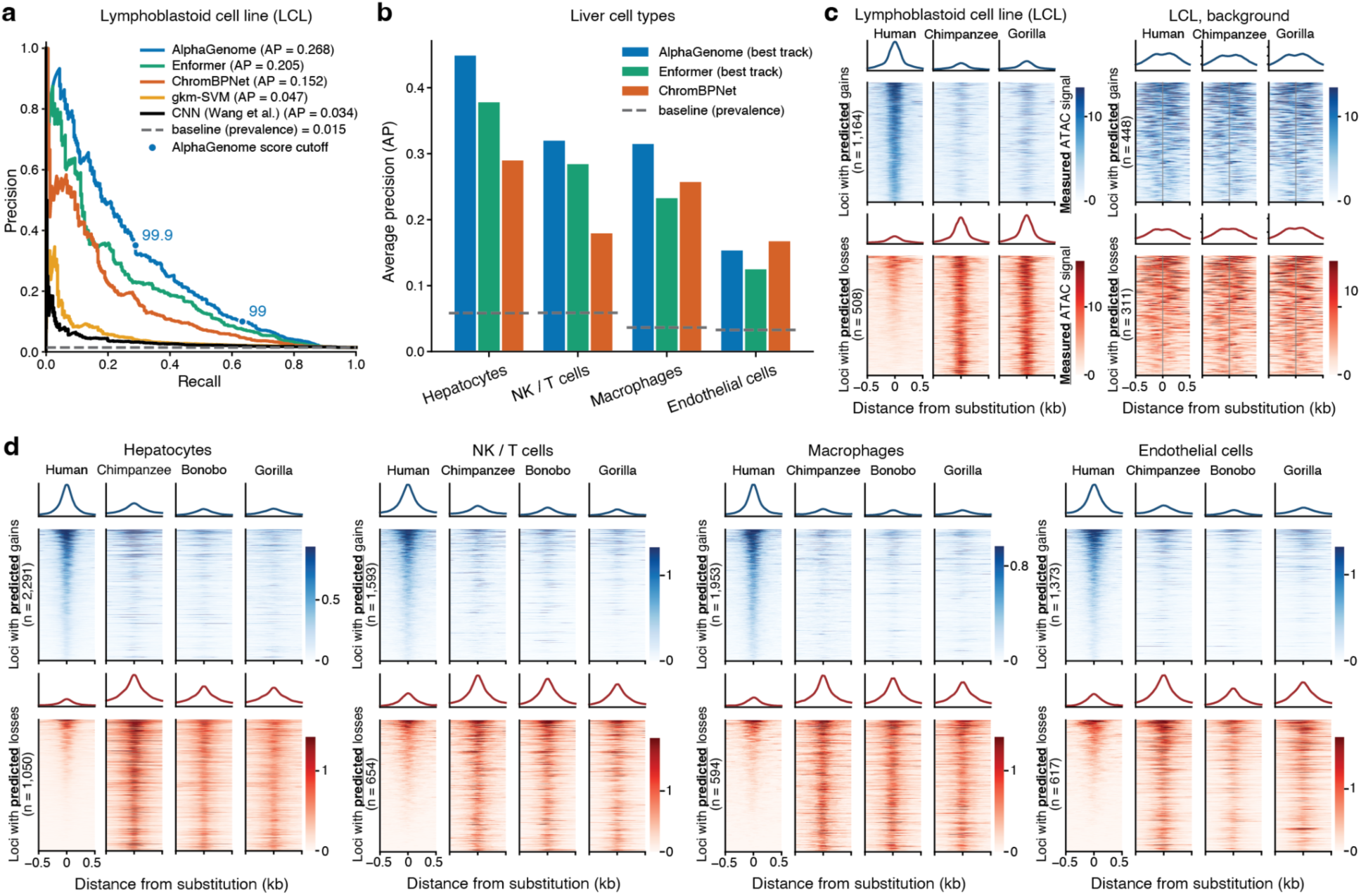
AlphaGenome predicts human-gained and human-lost cis-regulatory elements. **a**, Precision–recall curves and average precision (AP) for five sequence models (AlphaGenome, Enformer, ChromBPNet, gkm-SVM, and a classification CNN from Wang et al. 2026^13^) discriminating human-lost differentially accessible ATAC-seq peaks (n = 322) from conserved, non-significant background peaks (n = 21,738) against chimpanzee and gorilla in lymphoblastoid cell lines (LCLs; García-Pérez et al., 2021^16^). Each region is scored by the maximum human-lost-oriented score over substitutions in the same region. Dashed line, prevalence of the positive class against background (0.015). Circles on the AlphaGenome curve mark AlphaGenome score thresholds of 99 and 99.9. **b**, AP for AlphaGenome, Enformer and ChromBPNet in the same task as (**a**) in four liver cell types, for peaks called lost against chimpanzee, bonobo and gorilla (hepatocytes, n = 1,206 lost and 40,156 background; NK/T cells, 288 and 9,531; macrophages, 316 and 17,048; endothelial cells, 344 and 20,653). AlphaGenome and Enformer use each cell type’s best-performing tissue or cell type. Dashed lines, per-cell-type prevalence (0.016 to 0.029). **c,d**, Mean profile and per-locus heatmap of measured ATAC signal in LCLs (**c**) and the four liver cell types (**d**) from human and each non-human great ape, in ±500 bp windows at the orthologous position of every human-specific substitution passing the 99.9th percentile AlphaGenome score, gained above (LCLs n = 1,259; hepatocytes 2,441, NK/T 1,593, macrophages 1,953, endothelial 1,373) and lost below (LCLs n = 536; hepatocytes 1,090, NK/T 654, macrophages 594, endothelial 617). For LCLs, the matched background follows (n = 448 on the gained side and 311 on the lost side), being substitutions with near-zero AlphaGenome scores that pass the same ancestral-allele, peak-overlap, liftover and mappability filters as the called sets and constituting the complete eligible set rather than a subsample. All blocks are ascertained in the tier where the lost set still has positives, chimpanzee and gorilla for LCLs and chimpanzee, bonobo and gorilla for liver. Each species’ coverage is scaled by a single multiplicative factor equalizing strong, shared, non-differential peaks to human, so signal heights are comparable across species; the gained and lost blocks carry independent color scales and profile limits and are not comparable with each other, while each background block is drawn on the scale of the block it controls and values exceeding that ceiling are clipped. Loci ordered by descending human mean signal.

**Extended Data Fig. 2:**
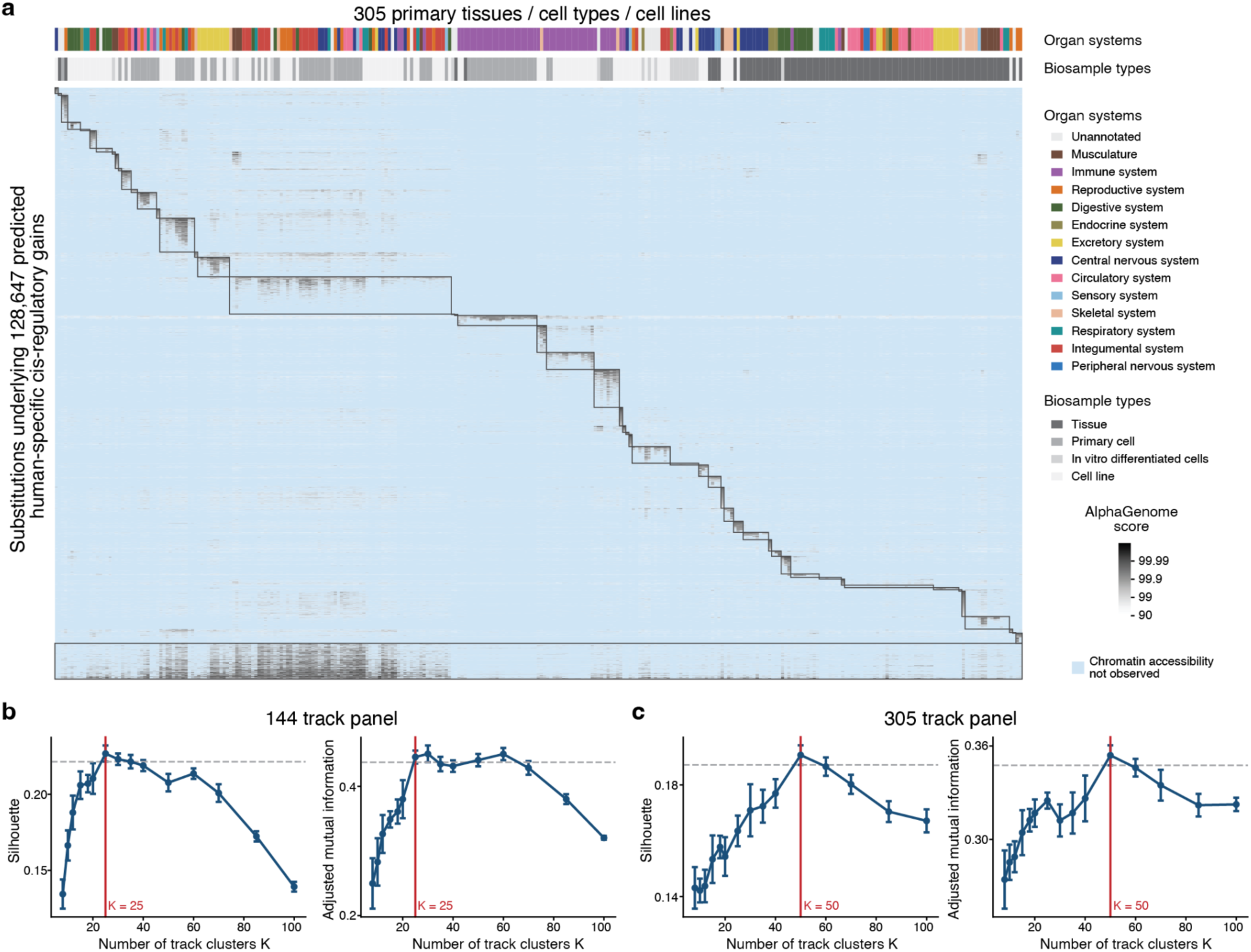
Clustering of predicted gained CREs based on tissue specificity. **a**, Clustered heatmap of predicted human-gained CREs across a 305-tissue panel of tissues, cell types, and cell lines. Rows are a random sample of 10,000 of the 128,646 open-chromatin elements called gained in at least one panel tissue or cell type at the 99.9th percentile, and columns are the 305 tissues or cell types, clustered into 50 programs based on correlation and average linkage. Color is a signed log_10_ transform of the AlphaGenome score, taking each element’s strongest substitution among those open in that tissue or cell type, and is shown only where the element lies in that tissue’s own open chromatin. **b,c**, Selecting the number of clusters for panels of 144 tissues (**b**) and 305 tissues (**c**) by silhouette score (left) and adjusted mutual information against ENCODE organ annotations (right). Both criteria are bootstrapped over the substitutions, and the chosen K is the smallest whose score lies within one standard error of the best, giving K = 25 for the 144-tissue panel and K = 50 for the 305-tissue panel.

**Extended Data Fig. 3:**
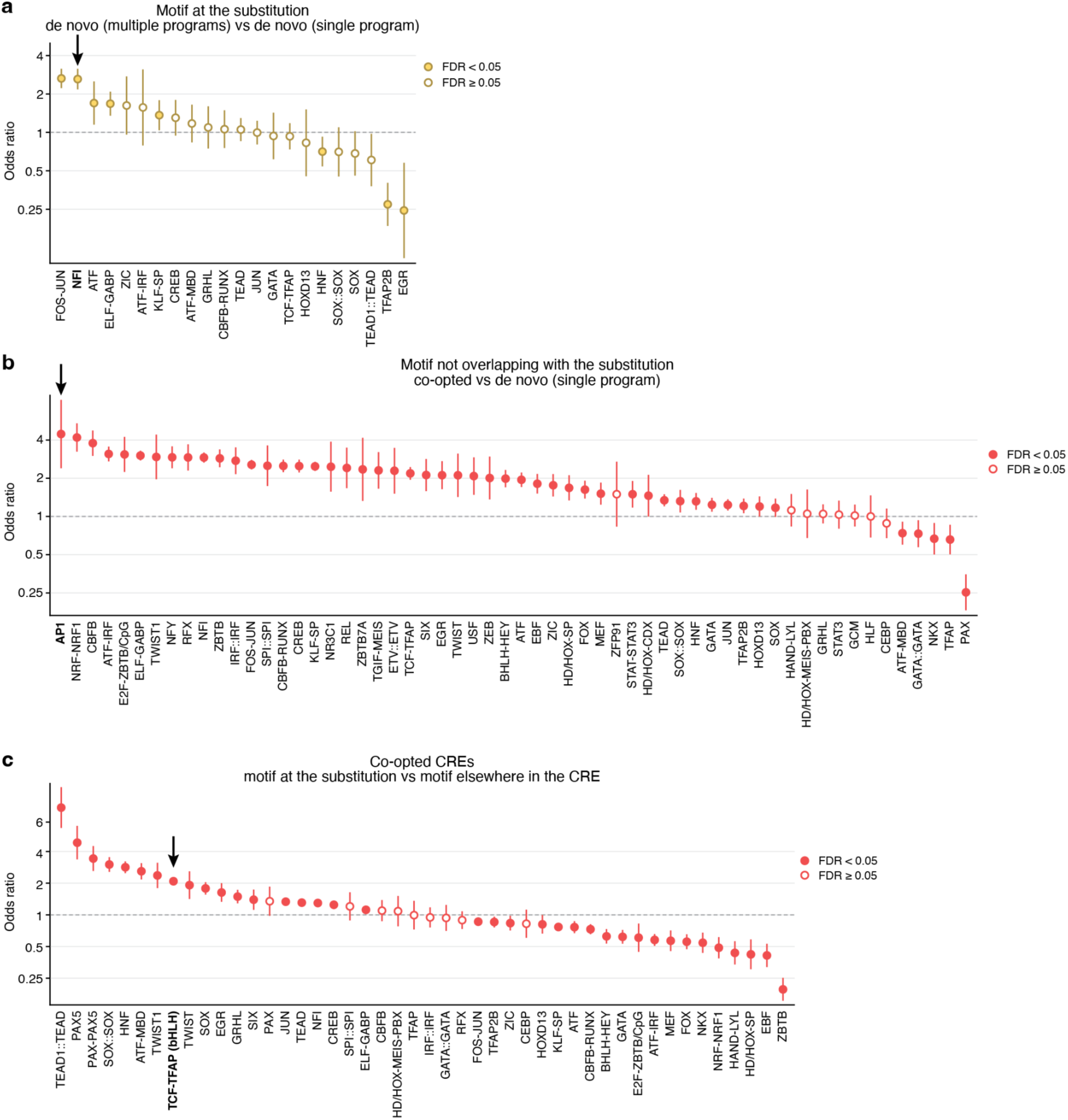
TF motif families in de novo and co-opted accessibility gains. Odds ratios for motif families among regions with human-specific cis-regulatory gains classified by mode of gain (Fig. 2b), with families ordered by odds ratio within each panel. **a**, Motif instances at the substitution, comparing de novo gains shared across programs with de novo gains confined to one program, for the 18 TF motif families with at least 50 instances at the substitution. **b**, Motif instances elsewhere in the CRE, outside the families present at the substitution, comparing co-opted CREs with de novo gains confined to one program, for the 59 TF families present in at least 200 CREs. **c**, Within co-opted CREs, the share of motif instances at the substitution against the share elsewhere in the same CREs, for the 40 TF families with at least 50 instances at the substitution. Values above 1 indicate families concentrated where the gain occurred. Points show Mantel–Haenszel odds ratios pooled over tissue programs with 95% confidence intervals. Filled points pass a Benjamini–Hochberg false discovery rate below 0.05 within the panel, and open points do not. TF families highlighted in the main text are indicated by arrows.

**Extended Data Fig. 4:**
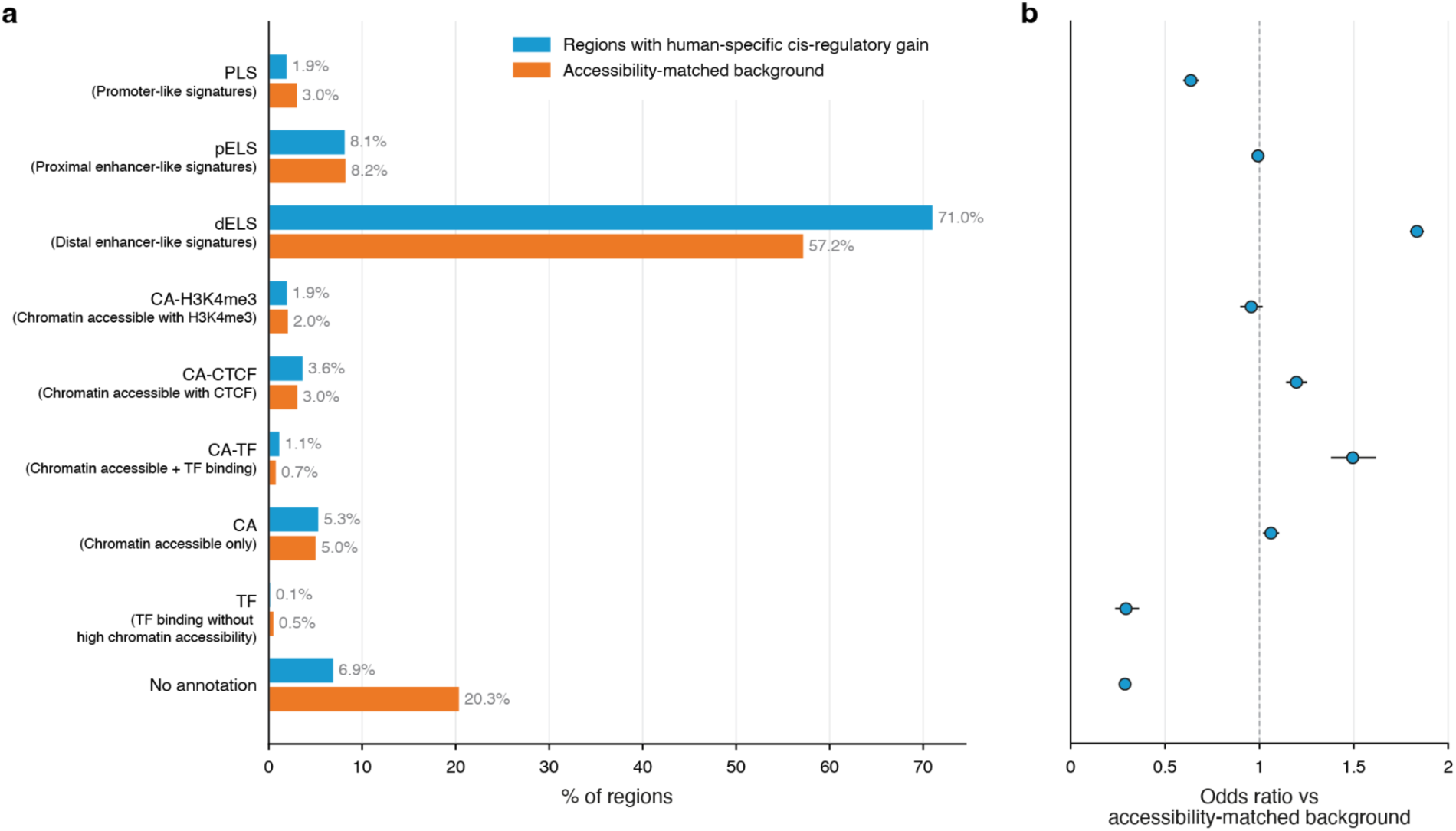
ENCODE candidate CRE (cCRE) classes of regions with predicted human-specific cis-regulatory gains. **a**, Proportion of regions in each class of the ENCODE candidate cis-regulatory element (cCRE) registry (v4, aggregated across biosamples), for the 71,327 regions with predicted human-specific cis-regulatory gains in the 144-track panel (excluding 18 regions on chromosome Y) and for accessibility-matched control regions. Each region is counted once and assigned to the cCRE containing one of its human-lineage substitutions, the gained substitution for predicted regions and one chosen at random for control regions. Control regions are the open-chromatin regions of the 144-track panel that carry at least one human-lineage substitution outside the predicted regions, reweighted to match the predicted regions in peak strength (the within-experiment signal percentile of the strongest overlapping DNase-seq peak across the 144 tracks) and peak width. **b**, Odds ratio of each class against all other classes, with 95% confidence intervals.

**Extended Data Fig. 5:**
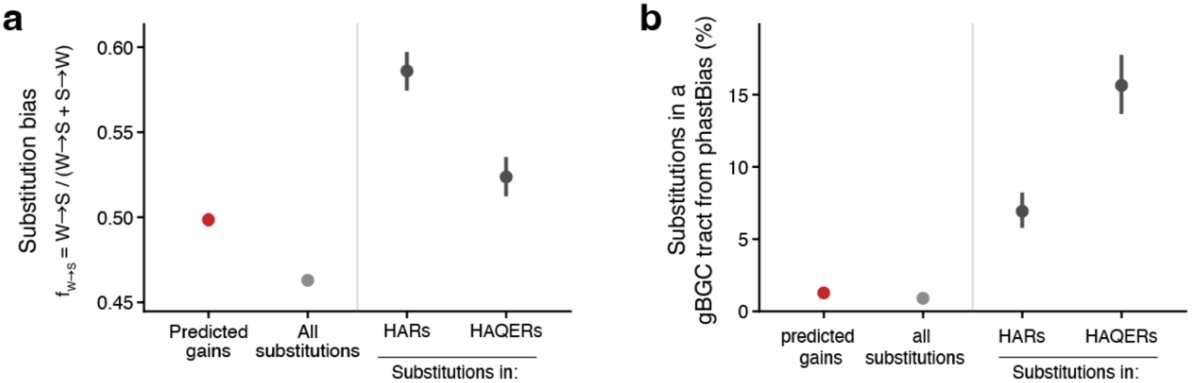
GC-biased gene conversion affects regions with predicted gains less than HARs and HAQERs. **a,b**, Substitution bias of human-lineage substitutions, f_W→S_ = (W→S) / (W→S + S→W), where W→S denotes weak-to-strong changes (A or T to G or C) and S→W the reverse. gBGC raises this ratio (**a**), and fraction of substitutions inside a gBGC tract called by phastBias (**b**). Sets are the substitutions with predicted gains in the 144-track panel (red), all human-lineage substitutions (gray), and HARs and HAQERs (black) for scale. Points are pooled estimates and bars are 95% block bootstrap intervals (1,000 resamples), resampling gained regions, HARs or HAQERs, or 1 Mb tiles for the genome-wide set.

**Extended Data Fig. 6:**
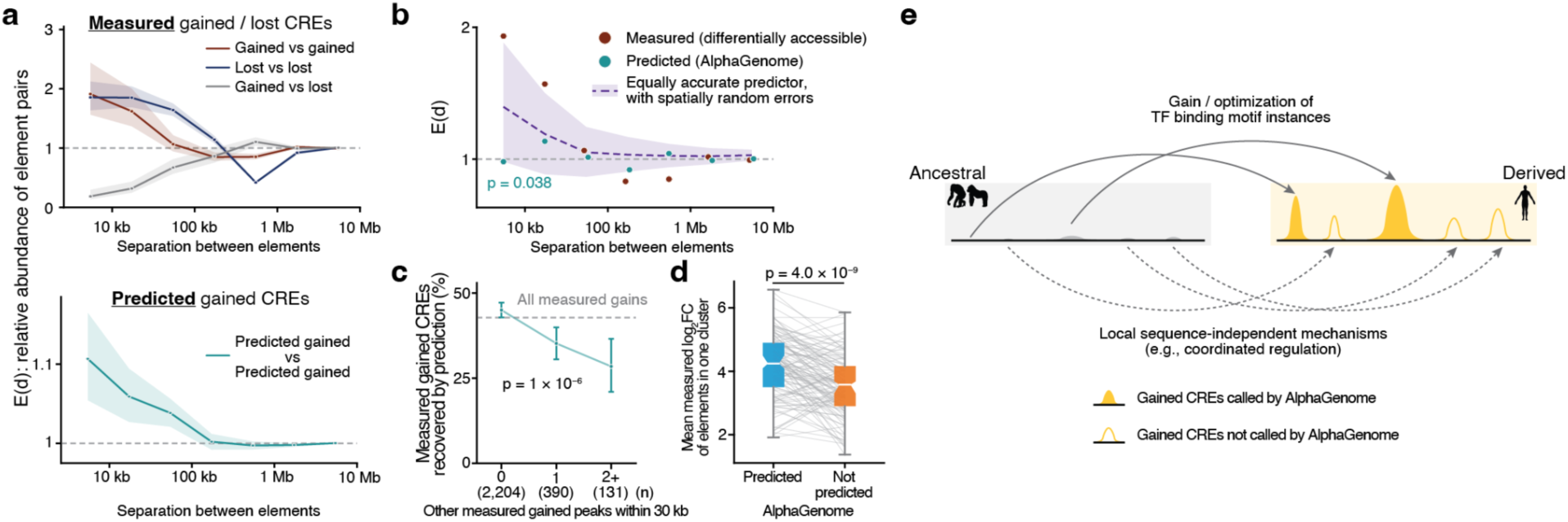
Predicted accessibility-gained CREs cluster spatially in the human genome, but less than measured accessibility-gained CREs. **a**, Pair enrichment E(d) was defined as the observed count of element pairs at separation d with both ends marked, over its mean under permutation. Element positions are held fixed and only the mark is permuted, within non-overlapping 1 Mb blocks. (Top) Measured differential accessibility in five cell types pooled (LCLs, hepatocytes, NK/T cells, macrophages, and endothelial cells; 2,725 gained and 2,476 lost peaks), decomposed into gained–gained, lost–lost and gained–lost. All three come from one permutation of a three-level label, preserving both marginal counts. (Bottom) Predicted gains of the same tissue program, with each program’s background restricted to the elements that the program opens. **b**, The controlled comparison. Measured and predicted gains scored on the same peaks, those the differential-accessibility test ran on, pooled over LCLs and liver hepatocytes, NK/T cells, macrophages and endothelial cells, with the predicted set sized to the measured one (n = 2,725 each) and the same binning and null for both, so the two arms differ only in what ranked the peaks. Points are unjoined and carry no intervals, because each bin is a separate comparison against the band rather than a point on a curve, and because a permutation interval would test a different null from the one the band poses. The purple band is a simulated predictor of the same accuracy and set size, its 1,166 correct calls drawn at random from the measured gains and its errors from the rest of the universe, which is what an imperfect but spatially unbiased detector would give. At 3 to 10 kb the predicted set holds 25 pairs against 100 measured and 36 expected under that band, one-sided *P* = 0.038 over 10,000 simulations. **c**, Proportion of measured human-gained peaks recovered by AlphaGenome, against the number of other measured gained peaks within 30 kb, over the five cell types with a paired table. Values are standardized over cell type (logistic on neighbor level plus cell type, averaged over the observed cell-type mix). Intervals are bootstrap resampled within cell type. The dashed line is the genome-wide recovery rate. **d**, Paired comparison, within each of 114 clusters of measured gained peaks (two or more within 20 kb, five cell types pooled), of the mean measured log_2_ fold change of the elements AlphaGenome calls against that of their uncalled neighbors. Boxes show the interquartile range, whiskers the 10th–90th percentiles, and notches the 95% confidence interval of the median **e**, Schematic of the systematic difference between measured and predicted accessibility-gained CREs within a cluster of measured accessibility-gained CREs.

**Extended Data Fig. 7:**
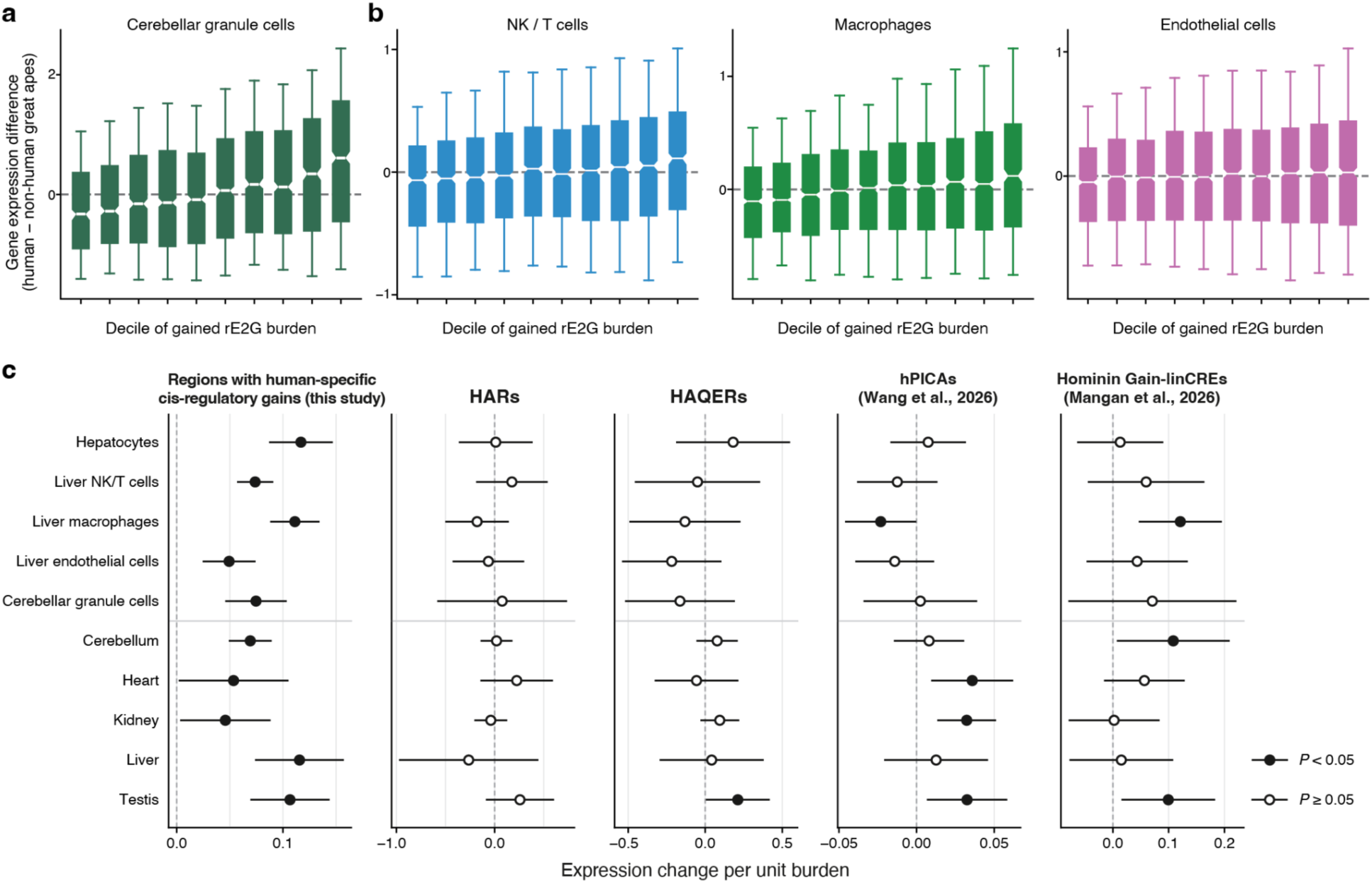
Association between gained rE2G burden and human-specific gene expression changes. **a**, Difference in gene expression between human and four non-human great apes (chimpanzee, bonobo, gorilla, orangutan) in NK/T cells, macrophages and endothelial cells, from the Yuan et al., 2025^18^ snRNA-seq pseudobulk, against deciles of rE2G burden of predicted gained CREs. Gained CREs are that cell type’s own snATAC peaks containing at least one substitution called human-gained by AlphaGenome, scored on the ENCODE biosample selected for that cell type (immature natural killer cell, suppressor macrophage and dermis blood vessel endothelial cell respectively), and rE2G burden is the sum of ENCODE-rE2G link scores from the same biosample over a gene’s gained CREs at the 99th percentile of the AlphaGenome score. Genes were retained above a symmetric expression floor on the average of the human and mean-ape values, leaving 13,563 genes in NK/T cells, 13,698 in macrophages and 13,186 in endothelial cells, giving 1,319 to 1,370 genes per decile. Expression differences are median-centered within the dataset. Boxes show the interquartile range, whiskers the 10th to 90th percentiles and notches the 95% confidence interval of the median. **b**, As in (**a**), difference in gene expression in cerebellar granule cells between human and bonobo, from the Sarropoulos et al. 2026^11^ snRNA-seq pseudobulk, against deciles of rE2G burden of predicted gained CREs. Gained calls are on the cerebellar cortex AlphaGenome track and links from the whole-cerebellum rE2G map, leaving 11,373 genes and 1,137 to 1,138 per decile. **c,** For each dataset, a CRE was counted as a candidate when it overlapped at least one region of the set, and a gene’s burden was the sum of ENCODE-rE2G link scores over its candidate CREs. Sets are CREs carrying a substitution at or beyond the 99th percentile of the AlphaGenome score (this study), human accelerated regions (HARs)^4,26^, human ancestor quickly evolved regions (HAQERs)^5^, human predicted increased chromatin accessibility regions (hPICAs)^13^, and hominin-lineage-gained CREs^14^. Rows are the cell types or tissues of gene expression datasets. Points show the coefficient of the burden on the human-versus-ape expression difference, adjusted for the burden of all other CREs linked to the gene, gene constraint (log_10_ s_het) and ancestral expression, with 95% confidence intervals. Filled points, *P* < 0.05.

**Extended Data Fig. 8:**
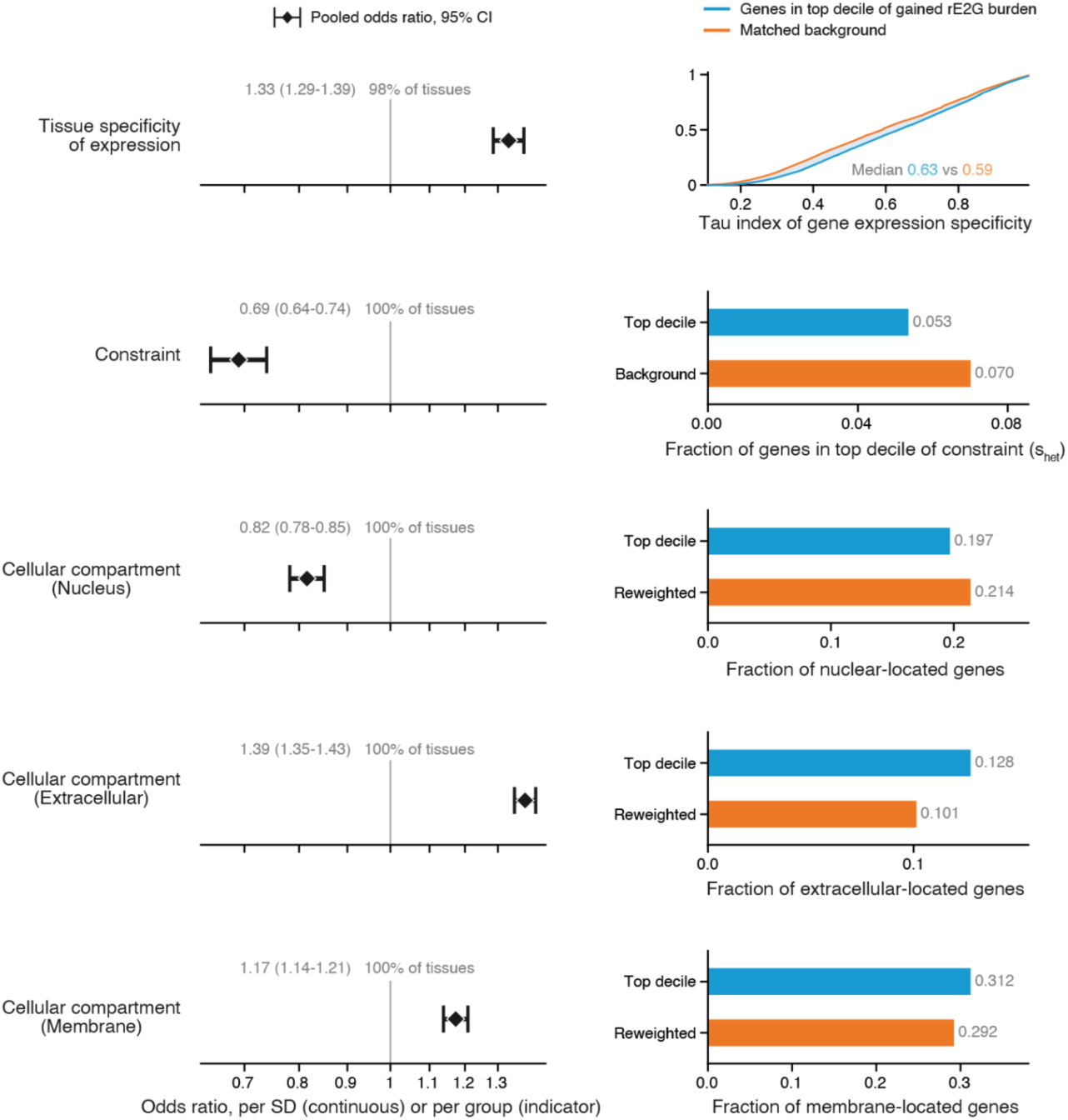
Gene properties associated with a high burden of predicted accessibility-gained CREs. Association of gene properties with the rE2G burden of predicted gained CREs, pooled over the 144 tissues or cell types. In each tissue, genes were ranked by rE2G burden, the sum of ENCODE-rE2G link scores over the gene’s CREs carrying at least one substitution past the 99th percentile of the AlphaGenome score in that tissue or cell type, and genes in the top decile were contrasted with the rest by logistic regression. The model included tissue specificity (tau over 51 human-protein-atlas consensus tissues), membership of the top decile of dosage sensitivity (s_het), and GO cellular-component annotation as nucleus, extracellular (secreted or extracellular matrix) or integral membrane. It also included regulatory complexity, the number of the gene’s links from CREs without a gained call, which is adjusted for but not shown. Continuous terms were z-scored within each tissue or cell type, all genes with at least one rE2G link and complete annotation were included. Left, odds ratio per standard deviation for tissue specificity and for group membership for the other properties, from a random-effects meta-analysis over tissues, with the 95% interval from a leave-one-cluster-out jackknife over the 25 tissue programs. The percentage is the fraction of tissues expected to share the direction of the pooled effect. Right, the same property in its own units for top-decile genes (blue) and for the remaining genes reweighted to match the top decile on every other property (orange), shown as cumulative distributions for tissue specificity and as proportions for the binary properties, each the median over tissues.

**Extended Data Fig. 9:**
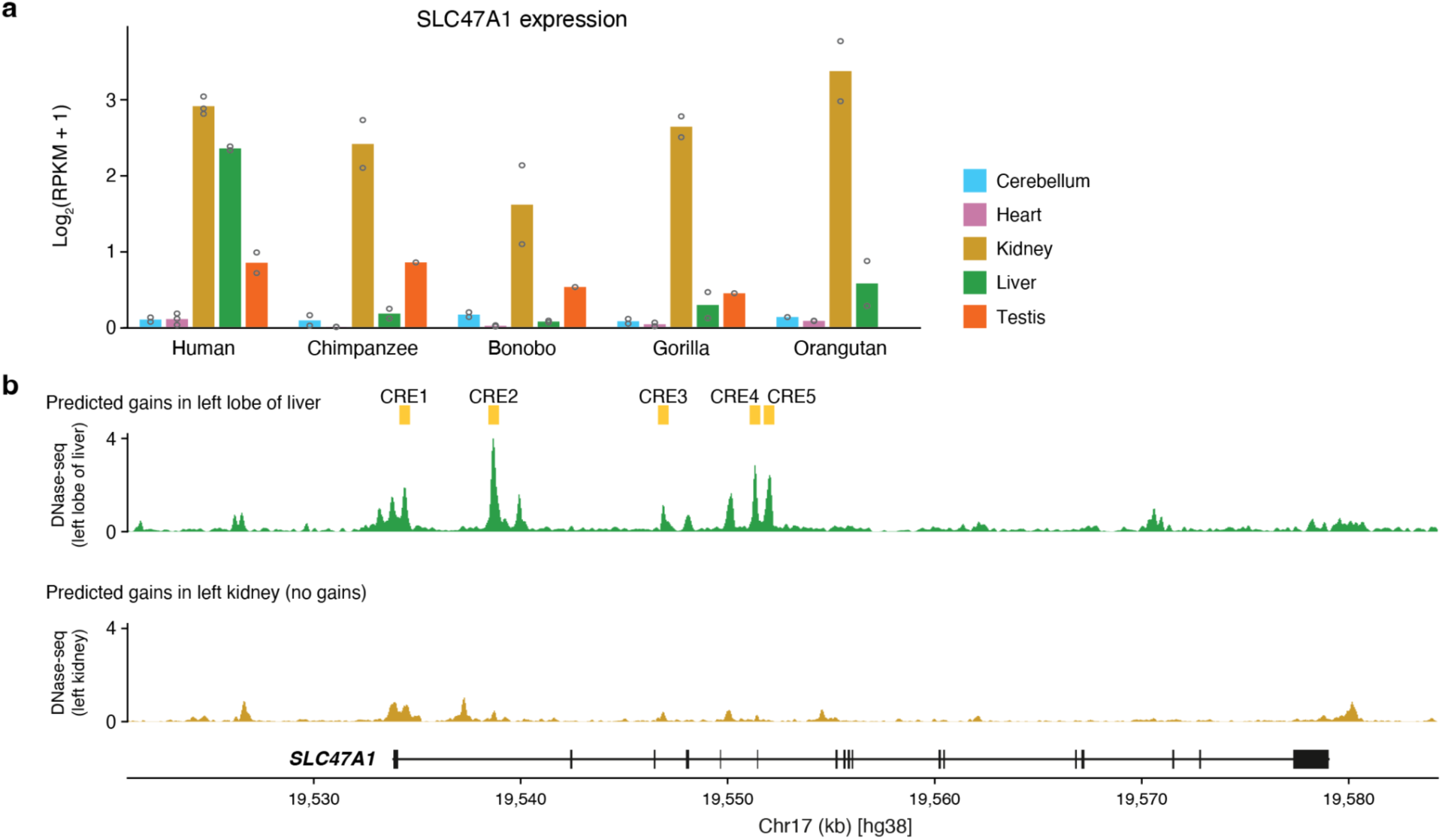
Liver-specific regulatory gain and expression increase of *SLC47A1* in the human lineage. **a**, SLC47A1 expression in five organs of human and four non-human great apes from bulk RNA-seq^30^, requantified against a single consensus gene annotation projected to every genome. Bars show the mean log_2_(RPKM + 1) for each species and organ, colored by organ, and open circles individual libraries. No orangutan testis library is available. **b**, Predicted human-gained CREs and measured DNase-seq accessibility at the *SLC47A1* locus, in the window of Fig. 3d (chr17:19,520,997–19,584,310), for left lobe of liver and left kidney. For each biosample, gained CREs are its ENCODE DNase peaks containing at least one substitution at or beyond the 99th percentile of that biosample’s AlphaGenome DNase score. Tracks show ENCODE DNase-seq signal for the same biosample, averaged over 50 bp bins and drawn on a shared y axis. No gained CRE is called in left kidney, where only one DNase peak is called in the window.

**Extended Data Fig. 10:**
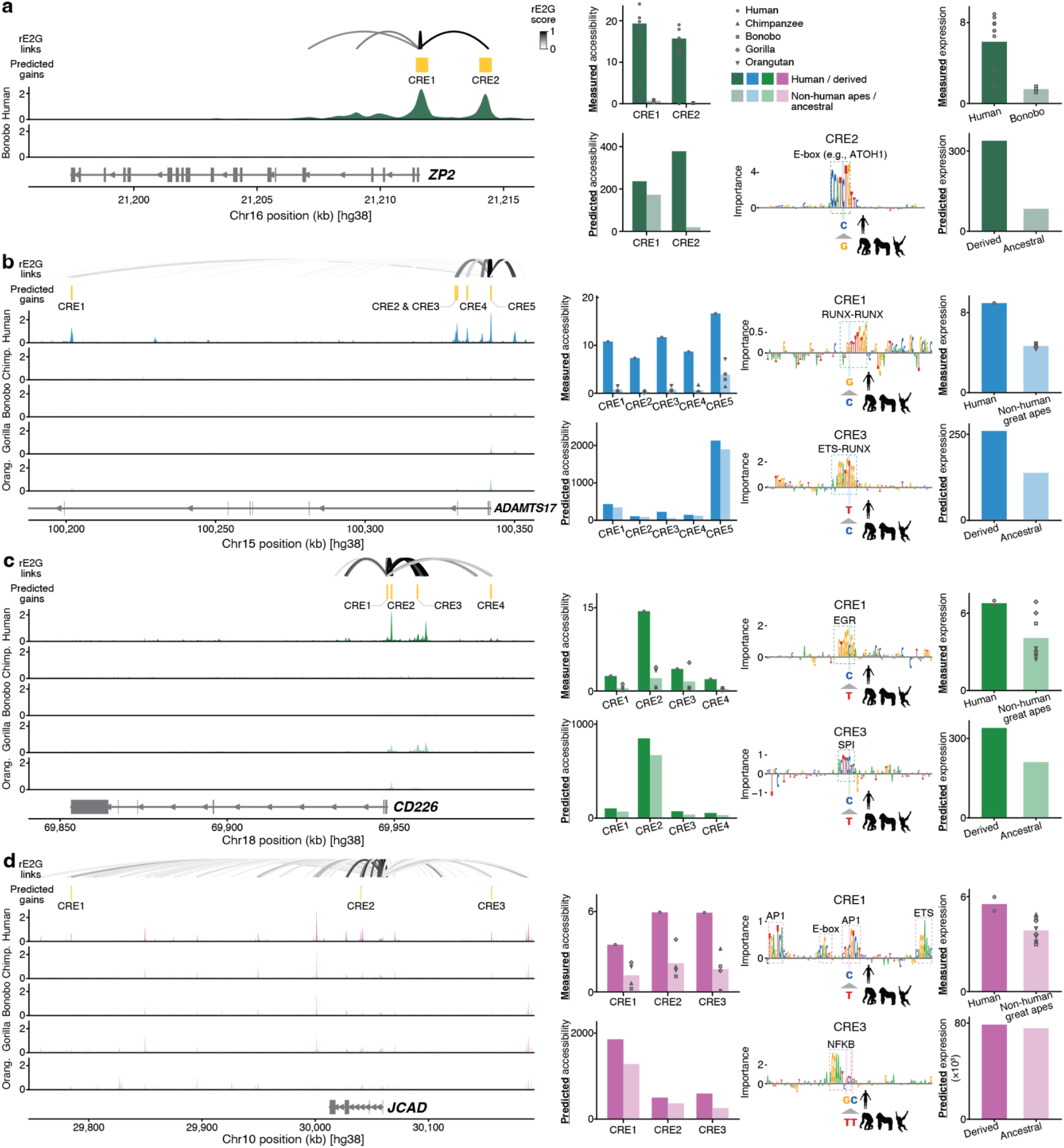
Predicted accessibility-gained CREs link human-specific substitutions to gene expression evolution in liver cell types and cerebellar granule cells. **a–d**, The *ZP2* (cerebellar granule cells; **a**), ADAMTS17 (NK/T cells; **b**), CD226 (macrophages; **c**), and *JCAD* (endothelial cells; **d**) loci. For each locus, from top: ENCODE-rE2G links to the gene’s TSS colored by link score; the predicted accessibility-gained CREs, two for *ZP2*, five for *ADAMTS17*, four for *CD226*, and three for *JCAD*; measured accessibility in human; measured accessibility in the non-human great apes at the orthologous positions; and the gene model. For each CRE, measured ATAC coverage in human against the great-ape mean, and AlphaGenome-predicted accessibility for the human sequence against the same sequence carrying the reconstructed ancestral alleles, on the biosample selected for that cell type. Contribution of each base of the human sequence to the AlphaGenome accessibility prediction, from in silico saturation mutagenesis over ±40 bp around each human-specific substitution, with high-importance matches to known TF binding motifs and the substitutions themselves annotated. Finally, measured expression of the gene in human and the non-human great apes by snRNA-seq, with points as donor pseudobulks, and AlphaGenome-predicted expression for the human sequence against the same sequence with all of the locus’s gained CREs reverted to their ancestral alleles simultaneously. The cerebellar panel (**a**) compares human with bonobo alone, the only outgroup in that dataset.

**Extended Data Fig. 11:**
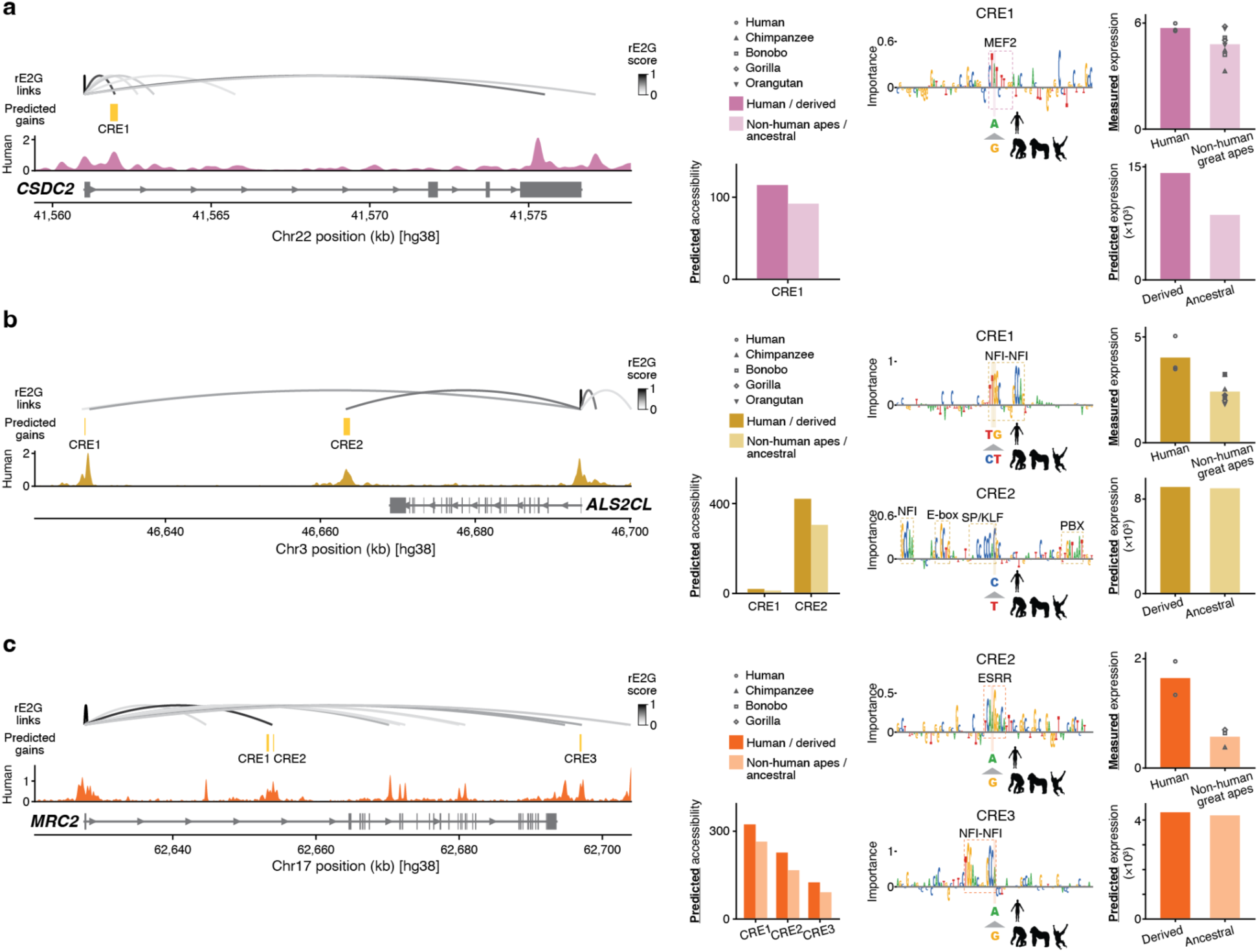
Predicted accessibility-gained CREs link human-specific substitutions to gene expression evolution in heart, kidney, and testis. **a–c**, The *CSDC2* (heart; **a**), *ALS2CL* (kidney; **b**) and *MRC2* (testis; **c**) loci. Unlike the loci in Fig. 3 and Extended Data Fig. 10, no cross-species accessibility data exist for these tissues, so the accessibility panels are human measurement against model prediction rather than human against ape, and the great-ape comparison enters only through bulk RNA-seq. For each locus, from top: ENCODE-rE2G links to the gene’s TSS colored by link score; the predicted accessibility-gained CREs, one for *CSDC2*, two for *ALS2CL* and three for *MRC2*; measured ENCODE DNase accessibility in human; and the gene model. For each CRE, AlphaGenome-predicted accessibility for the human sequence against the same sequence carrying the reconstructed ancestral alleles, on the biosample the tissue was scored with (right ventricle myocardium inferior, right kidney and testis respectively). Contribution of each base of the human sequence to the AlphaGenome accessibility prediction, from in silico saturation mutagenesis over ±40 bp around each human-specific substitution, with high-importance matches to known TF binding motifs and the substitutions themselves annotated. Measured expression of the gene in human and four non-human great apes by bulk RNA-seq from Brawand et al., 2011^30^, and AlphaGenome-predicted expression for the human sequence against the same sequence with all of the locus’s gained CREs reverted to their ancestral alleles simultaneously, read on the matching RNA track.

**Extended Data Fig. 12:**
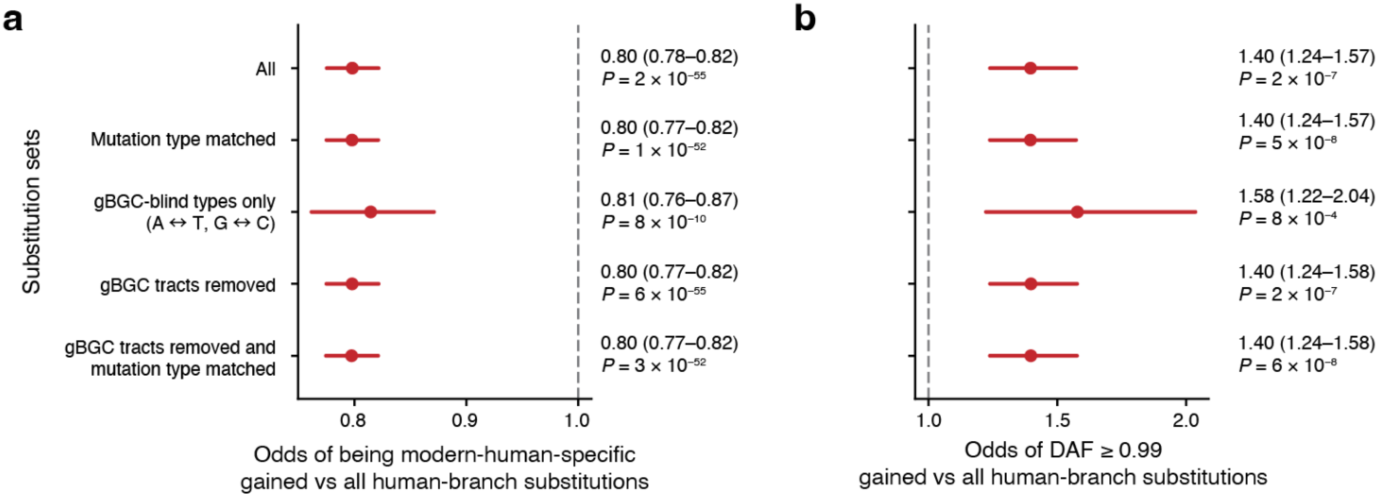
GC-biased gene conversion (gBGC) does not account for the age or the frequency of substitutions with predicted gains. **a,b**, Odds that a dated substitution with a predicted gain is modern-human-specific rather than shared with archaic hominins, relative to all human-lineage substitutions (**a**), and among modern-human-specific substitutions, odds of reaching a derived allele frequency of at least 0.99 in African populations, substitutions with gains relative to all human-lineage substitutions (**b**). Rows are the contrast itself, the same contrast stratified by mutation class, restricted to the two classes on which gBGC does not act (A↔T and G↔C), after removing substitutions inside a phastBias gBGC tract, and both corrections together. Points are odds ratios and bars 95% confidence intervals; dashed line, no difference.

## Supplementary Tables

**Supplementary Table 1**:

Peaks tested in the benchmark of human-specific chromatin accessibility change in LCLs and four liver cell types, with their labels and the human-lineage substitutions they contain.

**Supplementary Table 2**:

AlphaGenome and Enformer output tissue/cell type used in each benchmark comparison, with the performance instead based on a tissue fixed in advance by ontology match to the cell type.

**Supplementary Table 3**:

Regions with predicted human-specific cis-regulatory gains across 305 human tissues and cell types.

**Supplementary Table 4**:

Regions with predicted human-specific cis-regulatory gains across the 144 tissues and cell types used for clustering, with their tissue program assignments and genomic properties.

**Supplementary Table 5**:

Transcription factor binding motifs overlapping substitutions with accessibility-gained effects (AlphaGenome score ≥ 99.9) enriched among human-specific cis-regulatory gains in each tissue program.

**Supplementary Table 6**:

Gene Ontology (GO) terms enriched among genes near regions with human-specific cis-regulatory gains in each tissue program.

**Supplementary Table 7**:

Combinations of human CREs, rE2G maps and AlphaGenome predictions matched to each gene expression dataset.

**Supplementary Table 8**:

Genes in the top 10% by both gained rE2G burden and human-specific expression change in each dataset.

## References

1. King, M. C. & Wilson, A. C. Evolution at two levels in humans and chimpanzees. Science 188, 107–116 (1975).

2. Wray, G. A. The evolutionary significance of cis-regulatory mutations. Nat. Rev. Genet. 8, 206–216 (2007).

3. Avsec, Ž., et al. Advancing regulatory variant effect prediction with AlphaGenome. Nature 649, 1206–1218 (2026).

4. Pollard, K. S. et al. Forces shaping the fastest evolving regions in the human genome. PLoS Genet. 2, e168 (2006).

5. Mangan, R. J. et al. Adaptive sequence divergence forged new neurodevelopmental enhancers in humans. Cell 185, 4587–4603.e23 (2022).

6. Pääbo, S. The human condition-a molecular approach. Cell 157, 216–226 (2014).

7. Pollen, A. A., Kilik, U., Lowe, C. B. & Camp, J. G. Human-specific genetics: new tools to explore the molecular and cellular basis of human evolution. Nat. Rev. Genet. 24, 687–711 (2023).

8. Pampari, A. et al. ChromBPNet: bias factorized, base-resolution deep learning models of chromatin accessibility reveal cis-regulatory sequence syntax, transcription factor footprints and regulatory variants. bioRxivorg (2025) doi:10.1101/2024.12.25.630221.

9. Cheng, J. et al. AlphaGenome Atlas: in silico mutagenesis of the entire human genome improves prioritization and interpretation of non-coding variants. medRxiv 2026.09.16.26363192 (2026) doi:10.64898/2026.09.16.26363192.

10. Nagai, M., Murphy, A. E., Rizzo, K. & Koo, P. K. Toward generalizable and interpretable AI in regulatory genomics. Nat. Genet. 58, 1787–1802 (2026).

11. Sarropoulos, I. et al. The evolution of gene regulation in mammalian cerebellum development. Science 391, eadw9154 (2026).

12. Qiu, C. et al. Evolutionary transfer learning enables organism-wide inference of mammalian enhancer landscapes. bioRxivorg 2026.04.07.717039 (2026) doi:10.64898/2026.04.07.717039.

13. Wang, L., Li, Y., Han, D. & Wang, Z. Cross-species prediction reveals chromatin regions with increased accessibility in humans. Sci. Adv. 12, eady9169 (2026).

14. Mangan, R. J. et al. Sequence-to-function deep learning decodes human *cis* -regulatory evolution. bioRxiv 2026.08.24.746818 (2026) doi:10.64898/2026.08.24.746818.

15. Ferrando-Bernal, M. & Capra, J. A. Identifying human-specific transcription factor binding site gains and losses that contribute to human-specific phenotypes. bioRxiv 2026.09.06.747930 (2026) doi:10.64898/2026.09.06.747930.

16. García-Pérez, R. et al. Epigenomic profiling of primate lymphoblastoid cell lines reveals the evolutionary patterns of epigenetic activities in gene regulatory architectures. Nat. Commun. 12, 3116 (2021).

17. Avsec, Ž., et al. Effective gene expression prediction from sequence by integrating long-range interactions. Nat. Methods 18, 1196–1203 (2021).

18. Yuan, X., et al. The origin and molecular evolution of the mammalian liver cell architecture. bioRxiv (2025) doi:10.1101/2025.10.14.682049.

19. ENCODE Project Consortium. An integrated encyclopedia of DNA elements in the human genome. Nature 489, 57–74 (2012).

20. Yun, C. M., Ramalingam, V., Hecht, V. & Kundaje, A. ENCODE GRAMMAR: A deep learning model resource for decoding the DNA sequence logic of regulatory elements in the human genome. Genomics x AI Blog 10.5281/ZENODO.21795360 (2026) doi:10.5281/ZENODO.21795360.

21. Wu, X. et al. Evolutionary divergence in CTCF-mediated chromatin topology drives transcriptional innovation in humans. Nat. Commun. 16, 2941 (2025).

22. Prud’homme, B., Gompel, N. & Carroll, S. B. Emerging principles of regulatory evolution. Proc. Natl. Acad. Sci. U. S. A. 104 Suppl 1, 8605–8612 (2007).

23. McDonald, J. M. C. & Reed, R. D. Beyond modular enhancers: new questions in cis-regulatory evolution. Trends Ecol. Evol. 39, 1035–1046 (2024).

24. Liu, B. B. et al. Multiomics and deep learning dissect regulatory syntax in human development. Nature 653, 1240–1253 (2026).

25. Liu, Y. et al. Comparative single-cell multiome identifies evolutionary changes in neural progenitor cells during primate brain development. Dev. Cell 60, 414–428.e8 (2025).

26. Hubisz, M. J. & Pollard, K. S. Exploring the genesis and functions of Human Accelerated Regions sheds light on their role in human evolution. Curr. Opin. Genet. Dev. 29, 15–21 (2014).

27. Prescott, S. L. et al. Enhancer divergence and cis-regulatory evolution in the human and chimp neural crest. Cell 163, 68–83 (2015).

28. Boulanger, M. et al. Molecular co-accessibility identifies coordinated regulation between distant cis-regulatory elements. Mol. Cell 86, 3394–3410.e8 (2026).

29. Gschwind, A. R. et al. An encyclopedia of human enhancer-gene regulatory interactions. Nature 657, 179–190 (2026).

30. Brawand, D. et al. The evolution of gene expression levels in mammalian organs. Nature 478, 343–348 (2011).

31. Whalen, S. & Pollard, K. S. Enhancer function and evolutionary roles of human accelerated regions. Annu. Rev. Genet. 56, 423–439 (2022).

32. Kanton, S. et al. Organoid single-cell genomic atlas uncovers human-specific features of brain development. Nature 574, 418–422 (2019).

33. Kim, P. M., Korbel, J. O. & Gerstein, M. B. Positive selection at the protein network periphery: evaluation in terms of structural constraints and cellular context. Proc. Natl. Acad. Sci. U. S. A. 104, 20274–20279 (2007).

34. Huang, Y.-F. Dissecting genomic determinants of positive selection with an evolution-guided regression model. Mol. Biol. Evol. 39, (2022).

35. Otsuka, M. et al. A human transporter protein that mediates the final excretion step for toxic organic cations. Proc. Natl. Acad. Sci. U. S. A. 102, 17923–17928 (2005).

36. Staud, F., Cerveny, L., Ahmadimoghaddam, D. & Ceckova, M. Multidrug and toxin extrusion proteins (MATE/SLC47); role in pharmacokinetics. Int. J. Biochem. Cell Biol. 45, 2007–2011 (2013).

37. Florio, M. & Huttner, W. B. Neural progenitors, neurogenesis and the evolution of the neocortex. Development 141, 2182–2194 (2014).

38. Liu, R. & Jin, J.-P. Calponin isoforms CNN1, CNN2 and CNN3: Regulators for actin cytoskeleton functions in smooth muscle and non-muscle cells. Gene 585, 143–153 (2016).

39. Pollen, A. A. et al. Establishing Cerebral Organoids as Models of Human-Specific Brain Evolution. Cell 176, 743–756.e17 (2019).

40. Kim, S.-K. et al. Human-specific features of the cerebellum and ZP2-regulated synapse development. Cell 189, 1802–1819.e28 (2026).

41. Sepp, M. et al. Cellular development and evolution of the mammalian cerebellum. Nature 625, 788–796 (2024).

42. Prüfer, K. et al. The complete genome sequence of a Neanderthal from the Altai Mountains. Nature 505, 43–49 (2014).

43. Mafessoni, F. et al. A high-coverage Neandertal genome from Chagyrskaya Cave. Proc. Natl. Acad. Sci. U. S. A. 117, 15132–15136 (2020).

44. Prüfer, K. et al. A high-coverage Neandertal genome from Vindija Cave in Croatia. Science 358, 655–658 (2017).

45. Massilani, D. et al. A high-coverage Neandertal genome from the Altai Mountains reveals population structure among Neandertals. Proc. Natl. Acad. Sci. U. S. A. 123, e2534576123 (2026).

46. Meyer, M. et al. A high-coverage genome sequence from an archaic Denisovan individual. Science 338, 222–226 (2012).

47. Peyrégne, S., et al. A high-coverage genome from a 200,000-year-old Denisovan. bioRxiv (2025) doi:10.1101/2025.10.20.683404.

48. Zeberg, H., Jakobsson, M. & Pääbo, S. The genetic changes that shaped Neandertals, Denisovans, and modern humans. Cell 187, 1047–1058 (2024).

49. Moorjani, P., Amorim, C. E. G., Arndt, P. F. & Przeworski, M. Variation in the molecular clock of primates. Proc. Natl. Acad. Sci. U. S. A. 113, 10607–10612 (2016).

50. 1000 Genomes Project Consortium et al. A global reference for human genetic variation. Nature 526, 68–74 (2015).

51. Peyrégne, S., Boyle, M. J., Dannemann, M. & Prüfer, K. Detecting ancient positive selection in humans using extended lineage sorting. Genome Res. 27, 1563–1572 (2017).

52. Tambets, R. et al. Genetic analysis of circulating metabolic traits in 619,372 individuals. Nature 655, 971–978 (2026).

53. McCormick, D. B. & Chen, H. Update on interconversions of vitamin B-6 with its coenzyme. J. Nutr. 129, 325–327 (1999).

54. Murat, F. et al. The molecular evolution of spermatogenesis across mammals. Nature 613, 308–316 (2023).

55. Murphy, A., Nagai, M., Buendia, A., Kundaje, A. & Koo, P. Fine-tuning AlphaGenome in native JAX/Haiku. Genomics x AI Blog 10.5281/ZENODO.20272904 (2026) doi:10.5281/ZENODO.20272904.

56. Sheth, M. U. et al. Mapping enhancer-gene regulatory interactions from single-cell data. Nat. Genet. 58, 1941–1952 (2026).

57. Chen, S. et al. A genomic mutational constraint map using variation in 76,156 human genomes. Nature 625, 92–100 (2024).

58. Mallick, S. et al. The Allen Ancient DNA Resource (AADR) a curated compendium of ancient human genomes. Sci. Data 11, 182 (2024).

59. Kuderna, L. F. K. et al. Identification of constrained sequence elements across 239 primate genomes. Nature 625, 735–742 (2024).

60. Hubisz, M. J., Pollard, K. S. & Siepel, A. PHAST and RPHAST: phylogenetic analysis with space/time models. Brief. Bioinform. 12, 41–51 (2011).

61. Krueger, F. TrimGalore: Consistent Adapter and Quality Trimming for NGS, with Extra Functionality for RRBS Data. (Github).

62. Langmead, B. & Salzberg, S. L. Fast gapped-read alignment with Bowtie 2. Nat. Methods 9, 357–359 (2012).

63. Picard. https://broadinstitute.github.io/picard/.

64. Zhang, Y. et al. Model-based analysis of ChIP-Seq (MACS). Genome Biol. 9, R137 (2008).

65. Amemiya, H. M., Kundaje, A. & Boyle, A. P. The ENCODE Blacklist: Identification of Problematic Regions of the Genome. Sci. Rep. 9, 1–5 (2019).

66. Karimzadeh, M., Ernst, C., Kundaje, A. & Hoffman, M. M. Umap and Bismap: quantifying genome and methylome mappability. Nucleic Acids Res. 46, e120 (2018).

67. Dobin, A. et al. STAR: ultrafast universal RNA-seq aligner. Bioinformatics 29, 15–21 (2013).

68. Li, B. & Dewey, C. N. RSEM: accurate transcript quantification from RNA-Seq data with or without a reference genome. BMC Bioinformatics 12, 323 (2011).

69. Bray, N. L., Pimentel, H., Melsted, P. & Pachter, L. Near-optimal probabilistic RNA-seq quantification. Nat. Biotechnol. 34, 525–527 (2016).

70. Ma, S. et al. Molecular and cellular evolution of the primate dorsolateral prefrontal cortex. Science 377, eabo7257 (2022).

71. Francis Harrison, P. Varistran: Anscombe’s variance stabilizing transformation for RNA-seq gene expression data. J. Open Source Softw. 2, 257 (2017).

72. Li, H. et al. The Sequence Alignment/Map format and SAMtools. Bioinformatics 25, 2078–2079 (2009).

73. Quinlan, A. R. & Hall, I. M. BEDTools: a flexible suite of utilities for comparing genomic features. Bioinformatics 26, 841–842 (2010).

74. Robinson, M. D., McCarthy, D. J. & Smyth, G. K. edgeR: a Bioconductor package for differential expression analysis of digital gene expression data. Bioinformatics 26, 139–140 (2010).

75. Hansen, K. D., Irizarry, R. A. & Wu, Z. Removing technical variability in RNA-seq data using conditional quantile normalization. Biostatistics 13, 204–216 (2012).

76. Love, M. I., Huber, W. & Anders, S. Moderated estimation of fold change and dispersion for RNA-seq data with DESeq2. Genome Biol. 15, 550 (2014).

77. Zhu, A., Ibrahim, J. G. & Love, M. I. Heavy-tailed prior distributions for sequence count data: removing the noise and preserving large differences. Bioinformatics 35, 2084–2092 (2019).

78. Lee, D. et al. A method to predict the impact of regulatory variants from DNA sequence. Nat. Genet. 47, 955–961 (2015).

79. Lee, D. LS-GKM: a new gkm-SVM for large-scale datasets. Bioinformatics 32, 2196–2198 (2016).

80. Pedregosa, F. et al. Scikit-learn: Machine Learning in Python. arXiv [cs.LG] (2012) doi:10.48550/arXiv.1201.0490.

81. Johansen, N. J. et al. Evaluating methods for the prediction of cell-type-specific enhancers in the mammalian cortex. Cell Genom. 5, 100879 (2025).

82. Kempynck, N. et al. CREsted: modeling genomic and synthetic cell-type-specific enhancers across tissues and species. Nat. Methods 23, 946–959 (2026).

83. Lundberg, S. & Lee, S.-I. A unified approach to interpreting model predictions. arXiv [cs.AI] (2017) doi:10.48550/arXiv.1705.07874.

84. Shrikumar, A. et al. Technical Note on Transcription Factor Motif Discovery from Importance Scores (TF-MoDISco) version 0.5.6.5. arXiv [cs.LG] (2018).

85. Fi-NeMo: Identifying Transcription Factor Binding Sites from Neural Network Contribution Scores. (Github).

86. McLean, C. Y. et al. GREAT improves functional interpretation of cis-regulatory regions. Nat. Biotechnol. 28, 495–501 (2010).

87. Gu, Z. & Hübschmann, D. rGREAT: an R/bioconductor package for functional enrichment on genomic regions. Bioinformatics 39, btac745 (2023).

88. Siepel, A. et al. Evolutionarily conserved elements in vertebrate, insect, worm, and yeast genomes. Genome Res. 15, 1034–1050 (2005).

89. Burke, D. L., Ensor, J. & Riley, R. D. Meta-analysis using individual participant data: one-stage and two-stage approaches, and why they may differ. Stat. Med. 36, 855–875 (2017).

90. Paule, R. C. & Mandel, J. Consensus values and weighting factors. J. Res. Natl. Bur. Stand. (1977) 87, 377–385 (1982).

91. Robins, J. M., Hernán, M. A. & Brumback, B. Marginal structural models and causal inference in epidemiology. Epidemiology 11, 550–560 (2000).

92. Sato, T. & Matsuyama, Y. Marginal structural models as a tool for standardization. Epidemiology 14, 680–686 (2003).

93. ENCODE Project Consortium et al. Expanded encyclopaedias of DNA elements in the human and mouse genomes. Nature 583, 699–710 (2020).

94. Yanai, I. et al. Genome-wide midrange transcription profiles reveal expression level relationships in human tissue specification. Bioinformatics 21, 650–659 (2005).

95. Uhlén, M. et al. Proteomics. Tissue-based map of the human proteome. Science 347, 1260419 (2015).

96. Zeng, T., Spence, J. P., Mostafavi, H. & Pritchard, J. K. Bayesian estimation of gene constraint from an evolutionary model with gene features. Nat. Genet. 56, 1632–1643 (2024).

97. Ashburner, M. et al. Gene ontology: tool for the unification of biology. The Gene Ontology Consortium. Nat. Genet. 25, 25–29 (2000).

98. Gene Ontology Consortium. The Gene Ontology knowledgebase in 2026. Nucleic Acids Res. 54, D1779–D1792 (2026).

99. Gupta, S., Stamatoyannopoulos, J. A., Bailey, T. L. & Noble, W. S. Quantifying similarity between motifs. Genome Biol. 8, R24 (2007).

100. Duret, L. & Galtier, N. Biased gene conversion and the evolution of mammalian genomic landscapes. Annu. Rev. Genomics Hum. Genet. 10, 285–311 (2009).

101. Capra, J. A., Hubisz, M. J., Kostka, D., Pollard, K. S. & Siepel, A. A model-based analysis of GC-biased gene conversion in the human and chimpanzee genomes. PLoS Genet. 9, e1003684 (2013).

102. Zoonomia Consortium. A comparative genomics multitool for scientific discovery and conservation. Nature 587, 240–245 (2020).

